# Impaired mitochondrial stress signaling mediates bone loss in male mice in the absence of BNIP3

**DOI:** 10.64898/2026.04.06.710936

**Authors:** Li Tian, Victoria Van Berlo, Vivin Karthik, Joshua P Passarelli, Victoria E DeMambro, Patience Mudjgiwa, Calvin Vary, Anyonya R Guntur

**Affiliations:** Center for Molecular Medicine, MaineHealth Institute for Research, Scarborough, ME, USA; Tufts University School of Medicine, Tufts University, Boston, MA, USA; Graduate School of Biomedical Sciences and Engineering, University of Maine, Orono, ME, USA

**Author notes:** ***Corresponding author and reprint request should be addressed:*** Anyonya R Guntur PhD MaineHealth Institute for Research 81 Research Drive Scarborough Maine 04074. **Disclosure Statement:** All authors have no conflicting interests. Li Tian MD PhD: Laboratory of Endocrinology and Metabolism, Department of Endocrinology and Metabolism, West China Hospital, Sichuan University, Chengdu 610041, Sichuan, People’s Republic of China. Joshua P Passarelli BS: Tufts University School of Medicine.

**Keywords:** Mitophagy, Osteoblast, Glycolysis, Mitochondria, Oxidative phosphorylation

## Abstract

Osteoblasts generate bone by secreting collagen and mineralizing it in response to various signaling cues. We have previously shown that the majority of ATP generated by differentiated osteoblasts in response to glucose is through glycolysis in contrast to undifferentiated cells that are more dependent on oxidative phosphorylation. To confirm our previous findings, metabolomics was performed for unlabeled polar metabolites, revealing elevated glycolytic metabolites at the later stages of differentiation. Krebs cycle (TCA cycle) metabolites were also changed confirming metabolic rerouting with differentiation. We hypothesized that an increase in mitophagy shifts ATP generation towards glycolysis resulting in the observed bioenergetic and metabolic changes. Utilizing calvarial osteoblasts isolated from a mitophagy reporter mouse model (MitoQC), an increase in mitophagy and the mitophagy receptor, *Bnip3,* was observed with osteoblast differentiation. KD of *Bnip3* in osteoblasts inhibited differentiation and mineralization arising from impaired mitochondrial function. In vivo, male *Bnip3* null mice exhibited a significant decrease in osteoblast numbers resulting in lower bone mass. Mechanistically we identified decreased fusion and increased fission factors, impaired stress signaling and increased proapoptotic factors in the absence of *Bnip3*. These data demonstrate for the first time that BNIP3 expression and mitophagy during osteoblast differentiation are necessary for relieving mitochondrial stress to maintain optimal bone mass.

## Introduction

Mitochondria are central to coordinating energy metabolism through the generation of ATP by the electron transport chain and by regulating responses to substrate utilization, starvation and growth factor signaling^1^. As such, mitochondrial dysfunction has been implicated in bone loss with aging resulting in osteoporosis^2^. Studies have suggested that mesenchymal stem cells (MSCs) undergoing osteogenic differentiation have increased reliance on oxidative phosphorylation (OxPhos)^3, 4^, and data showing that inhibition of lactate dehydrogenase leads to increased bone mass supports this idea^5^. Furthermore, systematic characterization of the bioenergetic pathways utilized by MSCs, preosteoblasts, preadipocytes, and osteoblasts revealed that C3H/HeJ derived calvarial osteoblasts (COBs) are more oxidative than C57BL/6J COBs coinciding with the higher bone mass observed in C3H/HeJ mice^6^. In contrast, a number of reports suggest that differentiated osteoblasts rely more on aerobic glycolysis, to fuel increased energy needs for collagen production^7–9^. Our previous studies showed that differentiating osteoblasts utilize both OxPhos and glycolysis for generating ATP but at the terminal stage there is a shift towards glycolysis, with about 70% of ATP generated glycolytically^10^.

Metabolic reprogramming in osteoblasts may occur through an upregulation of mitophagy, similar to what has been observed in cardiac progenitor cells^11^. Manipulation of mitophagy is a mechanism that is co-opted for modulating and fine-tuning both glycolysis and OxPhos. Mitophagy is a process through which dysfunctional mitochondria are targeted to the lysosome for degradation and turnover. It occurs through two major pathways, the classical PINK1 (PTEN induced kinase 1) / PRKN (parkin RBR E3 ubiquitin protein ligase) ubiquitin-mediated pathway and through dedicated mitophagy receptors. The current identified mitophagy receptors include factors from the outer mitochondrial membrane, inner mitochondrial membrane, and mitochondrial matrix. BNIP3 (Bcl2/adenovirus E1B 19 kDa like interacting protein3), BNIP3L (Bcl2/adenovirus E1B 19 kDa like interacting protein3like, NIX), FUNDC1 (FUN14 domain containing 1), BCL2L13 (BCL2-Like 13) and FKBP8 are all outer mitochondrial membrane binding proteins. Known inner mitochondrial membrane mitophagy receptors include Prohibitin2 (PHB2) which interacts with PINK1/ PARKIN and Cardiolipin, a mitochondrial membrane phospholipid. A recently discovered mitochondrial matrix protein, NLRX1, (NOD-like receptor) also contains a LC3 interacting region (LIR) domain involved in mitophagy. One common aspect of these proteins is the utilization of LIR to target the tagged mitochondria for degradation. Triggering mitophagy results in recruitment of LC3 by mitophagy receptors through their LIR domains resulting in initiation and formation of the mitophagophore that is ultimately targeted for degradation in the lysosome. Other mitophagy related quality control processes include mitochondrial derived vesicles, piecemeal mitophagy and micro autophagy^12–14^. Recent studies using genetic tools to fluorescently tag mitochondria from osteoblasts has implicated mitochondrial fission related secretion of mitochondria as a mechanism for inducing osteogenic differentiation^15^.

Although previous studies have implicated the classical PINK1/ PRKN ubiquitin mediated pathway in regulating osteoblast function^16^, the role of mitophagy receptors and other related mechanisms in regulating osteoblast bioenergetics and differentiation is currently unknown. In this study, we targeted the mitophagy receptor BNIP3 to understand the role of mitophagy in regulating osteoblast differentiation. We report effects of 1) mitophagy during osteoblast differentiation using a novel mitophagy reporter transgenic mouse model, 2) effects of knockdown (KD) of BNIP3 in vitro on osteoblast transcriptome, proteome and bioenergetics, and 3) effects of global knockout of BNIP3 in vivo on bone. This is the first report where a mitophagy receptor is implicated in regulating osteoblast differentiation through bioenergetics, mitochondrial stress response, and cell survival.

## Results

### Metabolomic analysis of differentiating osteoblasts

To understand the metabolic landscape of differentiation in osteoblasts we isolated metabolites at Day7, Day14 and Day19 of differentiation and compared them to respective non differentiation controls. PCA analysis showed clear separation with days of differentiation with significant differences in various metabolic pathways **(Fig 1A).** This distinct clustering of differentiated versus non-differentiated MC3T3E1C4 cells, regardless of the timepoints, was confirmed through 3-D principal component analysis (PCA). Volcano plots for Day7, Day14 and Day19 comparisons **(Fig 1B-D)**. Notably, the 7-day differentiated, and non-differentiated groups were distinctly separated from the 14-day and 19-day groups, indicating a significant shift in the metabolic response beyond the 7-day time point. Individual increased and decreased metabolites shown in the volcano plots are plotted in more detail as waterfall plots comparing all three time points to their respective controls. These pathways include glycolysis, TCA cycle and related pathways (**SFig1-3**), as well as purine, pyrimidine and redox metabolism (**SFig4-6**). We identified that during osteoblast differentiation, glycolysis undergoes dynamic changes over time. On day 7, glycolysis is suppressed, accompanied by lower lactate levels. By Day14, glycolytic activity resumes, though lactate levels remain unchanged. On Day19, glycolysis is elevated, leading to increased lactate production **(SFig1)**. Concurrently, the tricarboxylic acid (TCA) cycle is suppressed in differentiated cells, several TCA intermediates, including succinate, malate, citrate, oxaloacetate, and isocitrate, are reduced, while 2-oxoglutarate and fumarate are elevated, suggesting metabolic rerouting rather than complete suppression **(SFig2)**. Beyond these shifts, broader metabolic changes occur during differentiation with changes in nucleotide and amino acid metabolism exhibiting increasing divergence over time, and glutathione depletion, characterized by decreased reduced glutathione and increased oxidized glutathione, suggests increasing oxidative stress. with osteoblast differentiation **(SFig6)**.

**Fig 1.**
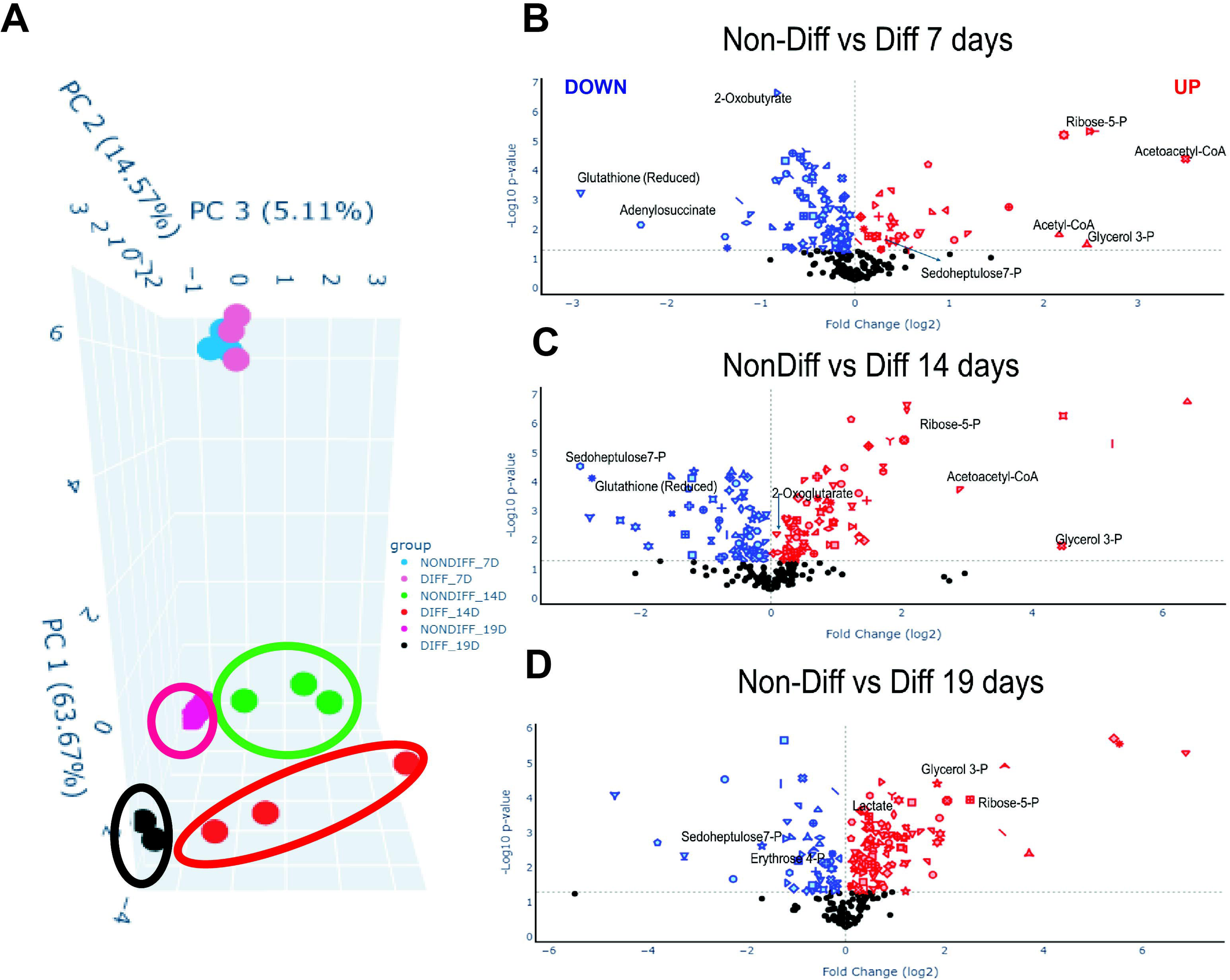
Temporal differentiation induces progressive molecular remodeling of the metabolome in Differentiating versus Non-differentiating osteoblasts. **(A)** Using an unlabeled polar metabolite profiling method, differentiating (DIFF) and non-differentiating (NONDIFF) samples formed unique clusters in a three-dimensional principal component analysis (PCA) of samples across all time points. Non-differentiating and differentiating groups segregate along PC1 (63.67%), while temporal progression from day 7 to day 19 separates primarily along PC2 (14.57%) and PC3 (5.11%). These patterns demonstrate a clear, time-dependent divergence in global metabolic profiles during the differentiation process. **(B-D)** Volcano plots showing differential abundance of detected metabolites between differentiating (DIFF) and non-differentiating (NONDIFF) osteoblasts at day 7 (B), day 14 (C), and day 19 (D). The x-axis represents log2 fold change (DIFF vs NONDIFF) and the y-axis represents –log10 p-value. Significantly increased features are shown in red, decreased features in blue, and non-significant features in black (dotted horizontal and vertical lines indicate statistical and fold-change thresholds, respectively). Progression from day 7 to day 19 reveals an increasing number and magnitude of significantly altered metabolites, consistent with progressive metabolic reprogramming during differentiation. Together, these data demonstrate that cellular differentiation is accompanied by progressive, coordinated metabolic remodeling, resulting in distinct and increasingly divergent molecular signatures over time. (n=3 biological samples each for all conditions, n=2 for differentiated 19D).

### Basal and induced mitophagy occurs in skeletal cells

To identify if any of the metabolic changes that were observed with differentiation are due to changes in mitochondrial turnover, we studied mitophagy. To identify if basal mitophagy occurs in osteoblasts and osteocytes we utilized the MitoQC transgenic mouse model. In this model, a mitochondrial target sequence from an outer membrane protein FIS1 was utilized to translate a tandem mCherry and GFP construct. Under basal conditions, mitochondria can be visualized using fluorescence microscopy in both red and green channels with the overlay being yellow. When mitophagy is activated and the mitochondria are engulfed by the lysosomes, green GFP signal is quenched and the mitochondria undergoing mitophagy show up as red punctae. We utilized wildtype MitoQC C57BL/6J newborn mouse hindlimbs to identify active mitophagy in articular chondrocytes, perichondrial cells, bone lining cells and osteocytes as observed in **(Fig 2A,B)**. Next, we treated primary calvarial osteoblasts isolated from homozygous MitoQC pups to induce mitophagy-using Deferiprone (1mM) an iron chelator. As seen in **(Fig 2C)** we see a significant increase in mitophagy (Red only punctae) with DFP treatment and quantified in **(Fig 2D)**. The increase in mitophagy correlated with an increase in BNIP3 protein levels after DFP treatment **(Fig 2E)**.

**Fig 2.**
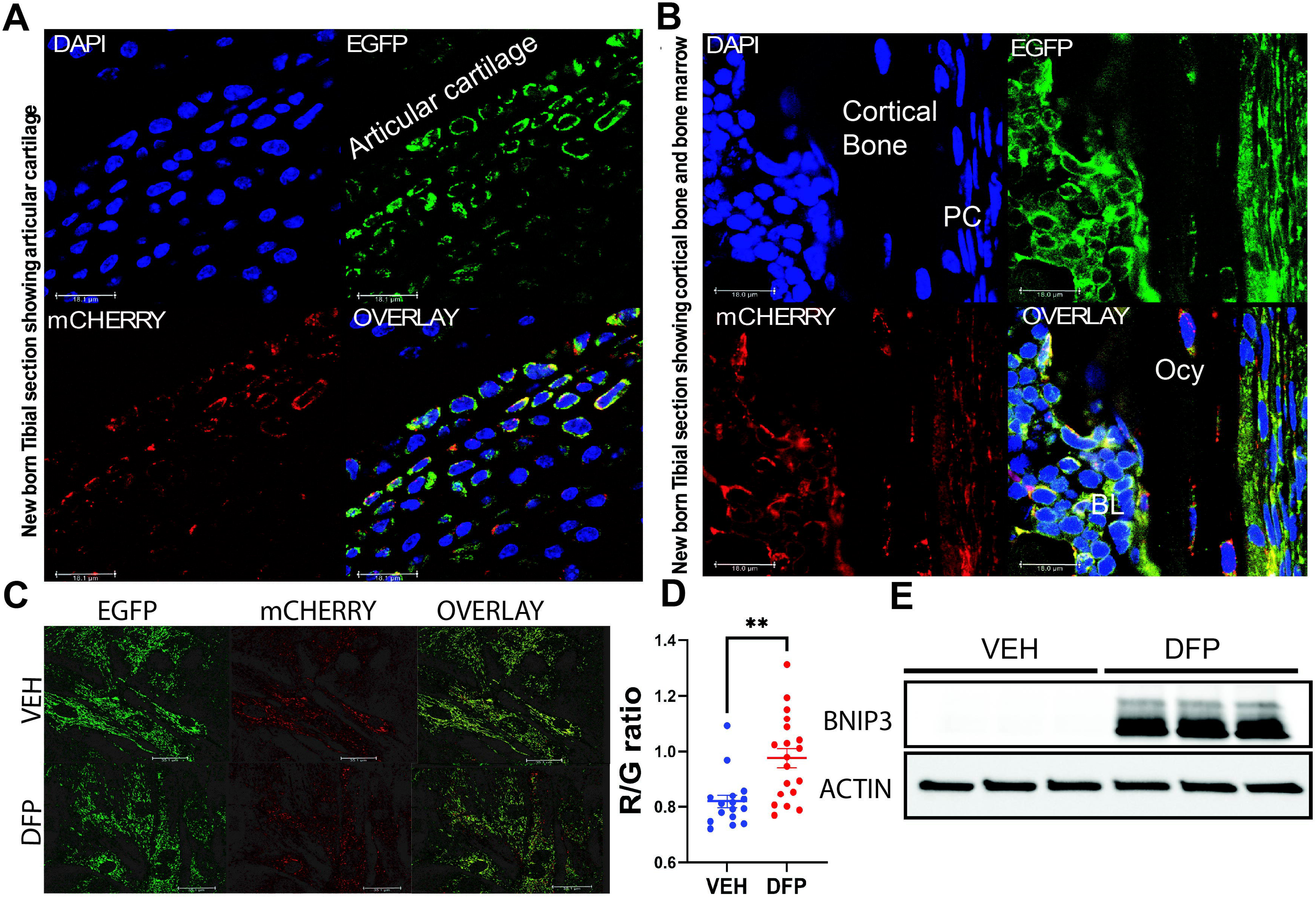
Mitophagy can be quantified in vivo in skeletal cells. **(A)** New born tibia from MitoQC mice show mitophagy (red punctae) in articular chondrocytes. **(B)** New born tibia from MitoQC mice show mitophagy (red punctae) in osteoblasts (near Bone lining (BL) and Perichondrium (PL) and osteocytes (Ocy). The different panels show DAPI stained nuclei, EGFP and MCherry indicate mitochondria and the overlay show red only punctae indicating mitophagy. **(C)** Calvarial osteoblasts (COB) from MitoQC mice treated with VEH (methanol, top pane) and DFP (1mM, bottom row). The red punctae increased with DFP treatment, indicating increased mitophagy. Representative confocal images from n=3 samples. **(D)** Quantification of Red/Green ratio from 16-20 cells using MitoQC counter (FIJI), comparing VEH to DFP. P-values from Student’s t-test (\*\**p* < 0.0001). (E) Western blotting results for BNIP3 and ACTIN after VEH or DFP treatment overnight in MC3T3E1C4 cells.

### Mitophagy increases during calvarial osteoblast differentiation and correlates with an increase in receptor mediated mitophagy

To identify if there is a physiological role for mitophagy during osteoblast differentiation we utilized calvarial osteoblasts isolated from MitoQC homozygous animals. We plated cells in glass covered tissue culture plates or cover slips and assessed mitophagy at various points from 0 days, 7 days, 14 days and 21 days post osteoblast differentiation and saw a significant increase in mitophagic punctae compared to predifferentiation. We observed and quantified a massive increase in mitophagy at 21 days of differentiation compared to Ndiff controls **(Fig 3A&B)**.We next utilized MC3T3E1C4 preosteoblast cells to identify factors that could potentially play a role in regulating mitophagy during osteoblast differentiation. We first identified *Bnip3* as being significantly induced with osteoblast differentiation in MC3T3E1C4 cells at the transcript level **(Fig 3C)**. We also confirmed this increase in *Bnip3* expression in primary calvarial osteoblasts after in vitro differentiation **(FigS7)** from a previously published RNAseq data set^17^. Next, we surveyed other mitophagy receptors such as *Bnip3L*, *Bcl2l213*, *Pink1* and saw a slight but significant increase in *Bnip3L* and *Bcl2l13* with a significant decrease in *Pink1* at the transcript level **(Fig 3C)**. We followed up these results in MC3T3E1C4 cells (preosteoblasts) to identify if BNIP3 colocalizes with mitochondria. Co-staining the cells with Mitotracker Red and BNIP3 antibody after treatment with cobalt chloride (CoCl_2_, 200µM), a pseudo hypoxia inducer, revealed a significant overlap of BNIP3 with mitochondria. Interestingly the mitochondria after overnight treatment with CoCl_2_ showed a profound change in shape assuming a doughnut like or rounded shape **(Fig 3D,E).** The effect of CoCl_2_ on mitochondrial structure translated into decreased OCR and an increase in ECAR when we performed a Mito stress test **(Fig 3F).** The increase seen in *Bnip3* also coincided with a significant increase in mitophagy **(Fig 3E).**

**Fig 3.**
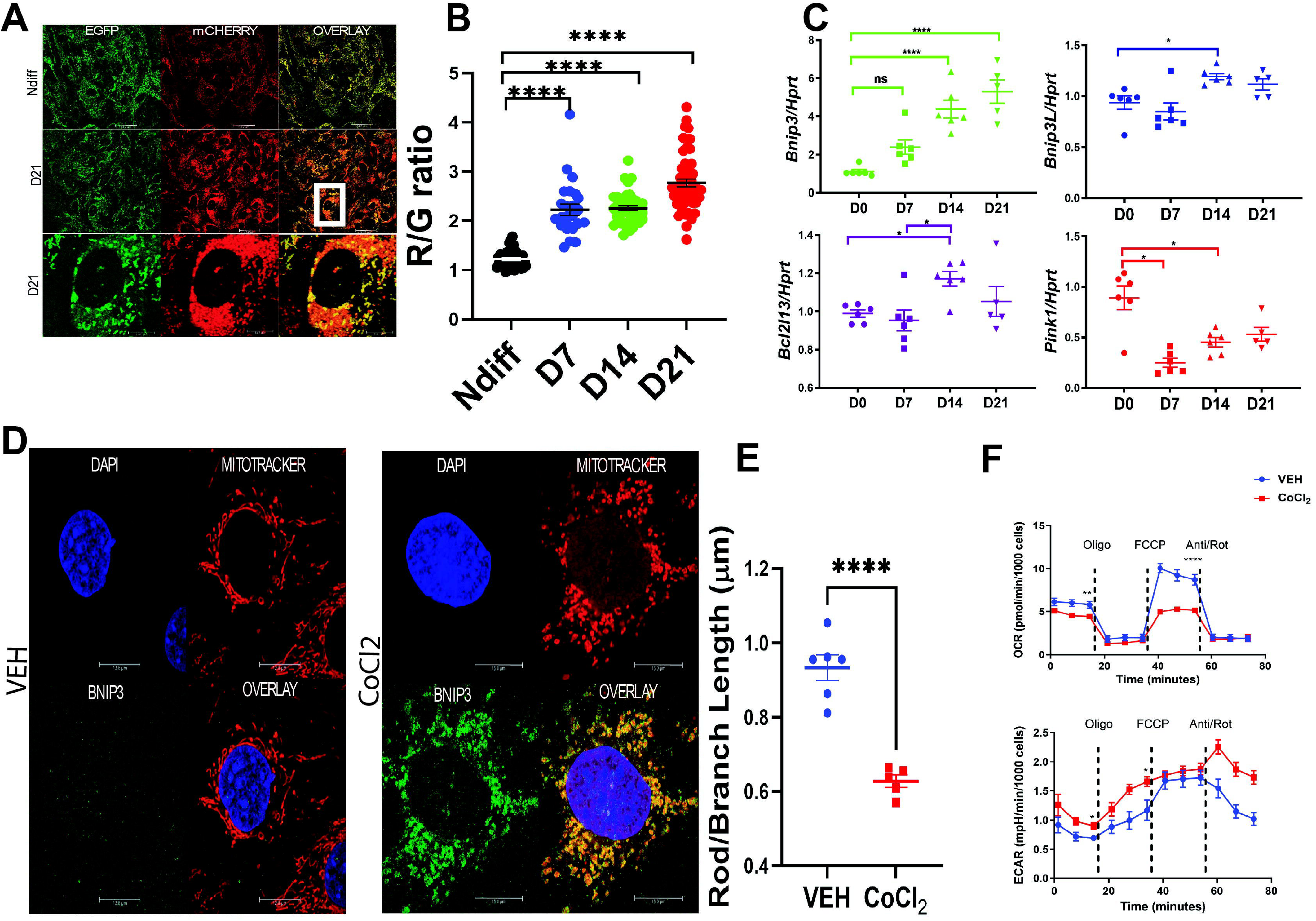
Mitophagy increases during osteoblast differentiation. **(A)** Calvarial osteoblasts (COB) from Nondiff (top) or 21 days in osteogenic media (DIFF, bottom panels). The red punctae indicate mitophagy. **(B)** Quantification of mitolysosomes/cell area from ≥30 cells using MitoQC counter (FIJI), comparing nondifferentiated cells to cells differentiated for 7, 14 or 21d, n=3 experiments. P-values from one-way ANOVA followed by Tukey’s post-hoc test (\*\*\*\**p* < 0.0001). **(C)** RT-qPCR on preosteoblasts compared to cells that have been differentiated for different timepoints D0 compared to D7, D14 and D21 of osteogenic differentiation including, *Bnip3*, *Bnip3l*, *Bcl2l13* and *Pink1*. P-values from one-way ANOVA followed by Tukey’s post-hoc test (*p < 0.05 **p < 0.01, ***p < 0.001 \*\*\*\**p* < 0.0001). (**D)** MC3T3E1C4 cells treated with VEH (left, top panel) and CoCl2 (200uM, Right panel). The different panels indicate DAPI (blue), FITC Green (BNIP3 staining), Mitotracker (Red) and overlay. Representative confocal images from n =3 experiments. **(E)** Quantification of Rod/branch lengths using MiRA plugin from ImageJ showing a significant decrease in networking after CoCl_2_ treatment. n=5-6 cells, P-values from students t-test (\*\**p* < 0.0001). **(F)** OCR (Top panel) and ECAR (bottom panel) after CoCl_2_ (red line); blue line is control (each with n>20 wells/group). There is a significant decrease in basal OCR with a compensatory increase in ECAR rates as indicated in the ECAR panel before oligomycin treatment. Asterisks indicate statistically significant P-values using Student’s t-test.

### Effects of knockdown of *Bnip3* on osteoblast differentiation with significant changes in the proteome of osteoblasts

The increase in *Bnip3* that we observed with osteoblast differentiation led us to next KD *Bnip3* in MC3T3E1C4 cells. We utilized adenoviral short hairpin KD to study the effect of reducing *Bnip3* expression during osteoblast differentiation. The KD when infected at a MOI of 100 was 70% at the transcript level, and BNIP3 protein was not detectable 3-days after KD **(Fig 4A)**. Upon osteoblast differentiation, *Bnip3* KD cells exhibited a significant decrease in terminal differentiation and mineralizing capacity **(Fig 4B,C)**. Significant decreases in expression of markers for both mitochondrial dynamics *Mfn2* and *Drp1* and osteoblast proliferation and differentiation including *Atf4*, *Osx* and *Bglap* **(Fig 4A)** were observed. To understand the changes that occur with differentiation at a global level we utilized Adeno scrambled and *Bnip3* KD cells to perform proteomic analysis on these cells using mass spectrometry. PCA analysis of the mass spectrometric data showed distinct clusters between the scrambled and *Bnip3* KD cells **(Fig 4D)**. We identified 3505 proteins using a cutoff threshold for p-values ≤0.05 and for enrichment fold changes ±Log2fold were used for a volcano plot. As shown in the analysis KD of BNIP3 showed significant upregulated and down regulated proteins in **(Fig 4E)**. These involved proteins from the inner mitochondrial membrane and mitochondrial ribosomes (OPA1, MRPL46), proteasome (PSMD14) and stress response (BCAT1, ASNS) among others indicating disruption in mitochondrial and other organellar homeostasis in osteoblasts. Taken together these data suggests that KD of *Bnip3* results in defects in osteoblast differentiation and mitochondrial metabolism.

**Fig 4.**
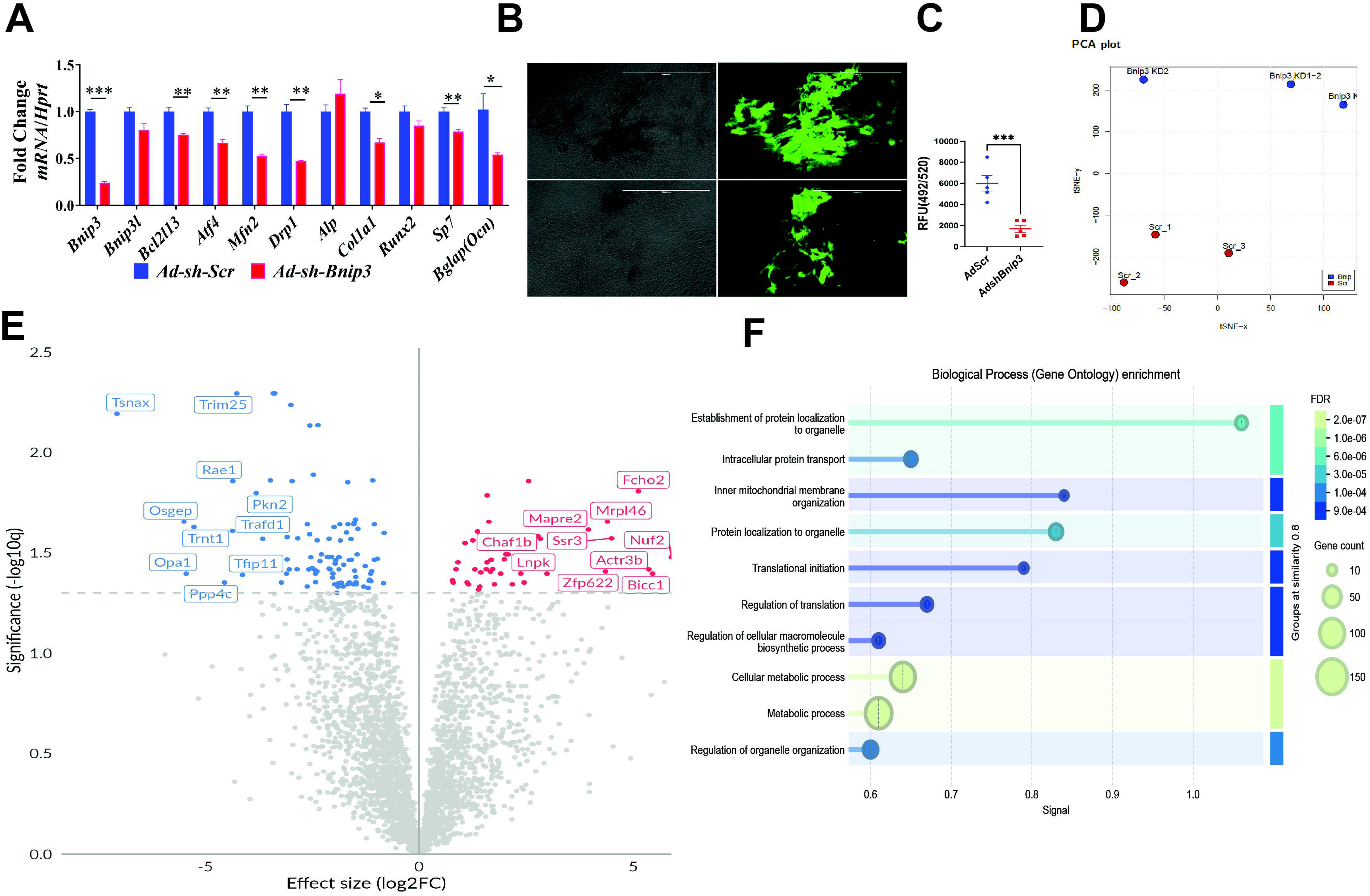
BNIP3 knockdown in vitro results in decreased osteoblast differentiation and changes in osteoblast proteome. **(A)** RT-qPCR of AdshRNAi *Bnip3* KD cells compared to scrambled control (n=3 replicates) on day 14 of differentiation BNIP3 KD significantly reduces expression of osteogenic and stress-response genes, including *Bnip3*, *Bnip3l*, *Bcl2l13*, *Atf4*, *Mfn2*, *Drp1*, *Alp*l, *Col1a1*, *Runx2*, *Sp7* and *Ocn*, normalized to *Hprt*. Asterisks indicate *p-*values from student’s t-test were significantly different. **(B)** Mineralization assay using an OsteoImage Mineralization Assay Lonza fluorescence assay kit comparing Scr control vs AdshRNAi *Bnip3* KD showing representative brightfield (left) and fluorescence (right) images. **(C)** Quantification of mineralization assay shown in B. Asterisks indicate P-values from Student’s t test (***p < 0.001). **(D)** PCA of proteomic data from AdshRNAi *Bnip3* KD and Scr controls cells. BigOmics Analytical software was utilized to perform principal component analysis resulting in separate clusters demonstrating a distinct proteomic shift driven by loss of BNIP3. Replicates cluster separately within their respective groups. **(E)** Volcano plot of differentially expressed proteins analyzed by mass spectrometry reveals significant up and down-regulated proteins (highlighted in red and blue, respectively) following BNIP3 KD. **(F)** Gene Ontology (GO) biological process enrichment analysis of differentially expressed genes reveals significant alterations in pathways related to protein localization to organelles, mitochondrial inner membrane organization, intracellular protein transport, translational initiation, regulation of translation, and cellular metabolic processes. Dot size reflects gene count and color indicates false discovery rate (FDR).

### Knockout of *Bnip3* shows a dimorphic bone phenotype with impaired bone mass in male mice

To study the effect of loss of *Bnip3* on bone mass and body composition in vivo, we performed a longitudinal DEXA analysis on both male and female mice at 8, 12 and 16 weeks of age. Female mice did not show any significant differences. Male mice exhibited a significant decrease in areal BMD, areal BMC, and femoral BMD, without any significant changes in body weight, lean mass, or fat mass **(FigS8)**. We next performed µCT analysis on femurs at 16 weeks of age to investigate the effect of loss of *Bnip3* on bone microarchitecture. We first performed µCT analysis on femoral metaphysis for trabecular bone. Female mice did not show any significant differences in trabecular or cortical bone microarchitecture and were similar to age matched controls. Male *Bnip3*^-/-^ mice on the other hand showed a significant decrease in a number of different trabecular parameters. BV/TV was significantly lower in the *Bnip3*^-/-^ femurs. There was also a significant decrease in BMD and trabecular thickness; there was no difference in connectivity density, trabecular number or spacing, suggesting that the phenotype could be because of the thinner trabeculae. There was a significant increase in SMI and bone surface/bone volume in the *Bnip3*^-/-^ femurs **(Fig 5A-C)**. Overall, there was a significant loss in trabecular bone in the absence of *Bnip3*. Next, we assessed cortical bone microarchitecture. Loss of *Bnip3* in male mice resulted in a significant decrease in cortical area and total area, cortical thickness, marrow area, cortical tissue mineral density cortical porosity did not reach significance. Polar moment of inertia, minimum and maximum moment of inertia were also significantly lower in the *Bnip3*^-/-^ males **(Fig 5B)**. These data suggest a sexually dimorphic bone phenotype in the absence of BNIP3 and the male mice have a significant bone phenotype with a decrease in bone mass. The bones based on the polar moment of inertia are weaker compared to WT controls. Collectively, these data show that deletion of *Bnip3* leads to trabecular bone loss, as well as a defect in cortical expansion in male mice while females are unaffected.

**Fig 5.**
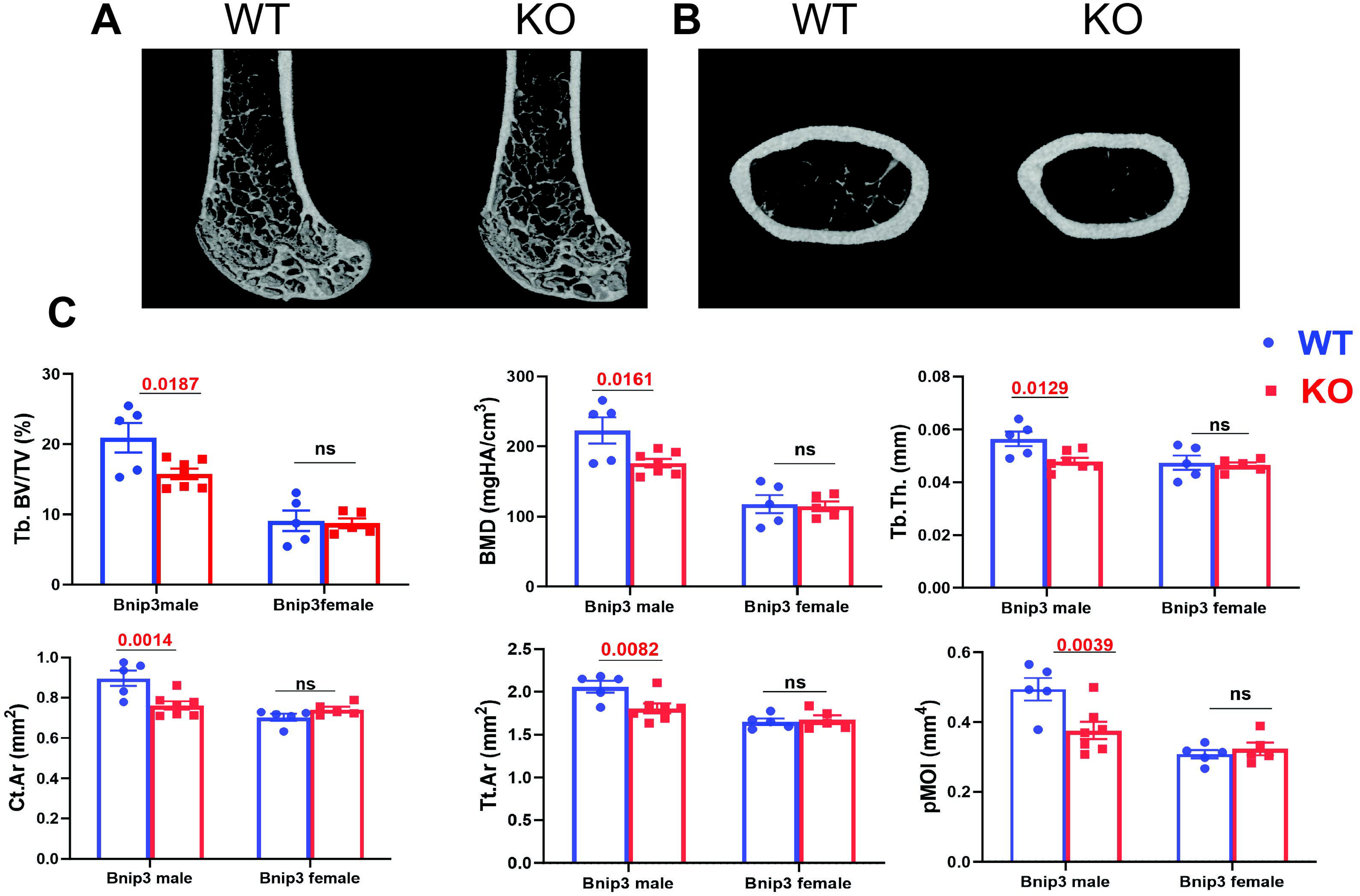
*Bnip3* deletion affects bone in male but not female mice. **(A,B)** Shown are representative μCT images of male femora showing trabecular and cortical bone. **(C)** Quantification of bone measurements comparing effects of sex (n=5-7 males, n=5 females) and genotype in 16 week old mice. Trabecular bone volume, trabecular bone mineral density, trabecular thickness, cortical area, total periosteal area, and polar moment of inertia were reduced in males lacking *Bnip3* compared to their wildtype littermates. P-values from 2 way-ANOVA are shown. ns = non-significant.

### Bone histomorphometry

Based on our µCT observations we next performed histologic analysis on undecalcified plastic embedded male 16-week-old femurs. A significant decrease in osteoblast number with no changes in osteoclast number was observed **(Fig 6A)**. Dynamic histomorphometry revealed significant decreases in MS/BS (mineral surface/bone surface) and MAR (mineral apposition rate). However, we could not compare BFR (bone formation rate) because a number of KO animals did not show a clear double label. In summary, *Bnip3^-/-^*mice had reduced bone density and impaired osteoblast numbers and activity. We confirmed decreased osteoid by staining undecalcified plastic embedded femur 5µm sections with trichrome staining **(Fig 6B)**. We did not find any significant changes in bone marrow adipocytes and osteoclasts (TRAP staining). Overall based on our *in vitro* and *in vivo* studies of loss of BNIP3 function studies we conclude that there is a significant defect in osteoblast numbers and function resulting in the low bone mass phenotype that is observed in the males.

**Fig 6.**
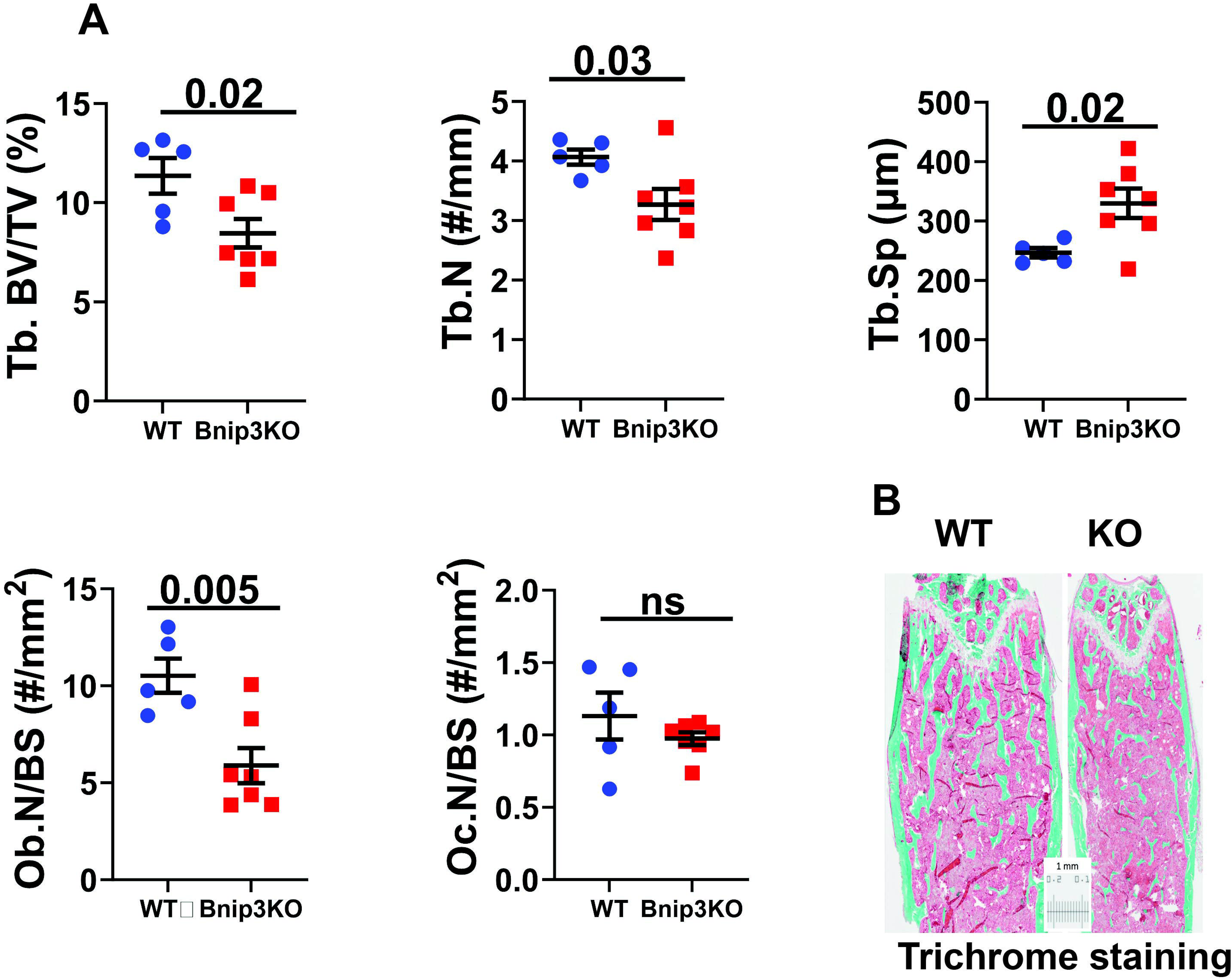
Histomorphometry analysis. **(A)** In WT and *Bnip3*^-/-^ males, trabecular %bone volume/total volume, trabecular number and trabecular spacing, osteoblast number per bone surface and osteoclast numbers per bone surface were quantified in the femur. Graphed are means ± SEM from 16 week old male mice, n=5-7 males, P-values from student’s t-test are shown. ns, not significant. **(B)** Representative image of trichrome staining of femora from WT and *Bnip3*^-/-^ mice. Images shown are representative of at least n=3 each.

### Loss of *Bnip3* impairs mitochondrial oxidative phosphorylation and function

*Bnip3* has been shown to regulate oxidative phosphorylation in adipocytes previously^18^. To determine whether the effects we observed are linked to metabolic changes, we examined mitochondrial function in control and *Bnip3* KD cells after 7 days of osteoblast differentiation. We performed a mitostress test to assess oxidative phosphorylation (OCR) and extracellular acidification rates. KD of *Bnip3* led to a reduction in several mitochondrial parameters compared to scrambled controls. Specifically, after FCCP treatment, we observed a decrease in spare respiratory capacity in *Bnip3* KD cells. While basal respiration was only slightly reduced, KD cells exhibited increased proton leak following Oligomycin treatment, suggesting uncoupling of oxidative phosphorylation from ATP generation **(Fig 7)**. Additionally, there was a significant increase in basal extracellular acidification rate (ECAR), indicating a metabolic shift toward glycolysis, likely as a compensatory response to impaired oxidative phosphorylation (**Fig 7A,B**).

**Fig 7.**
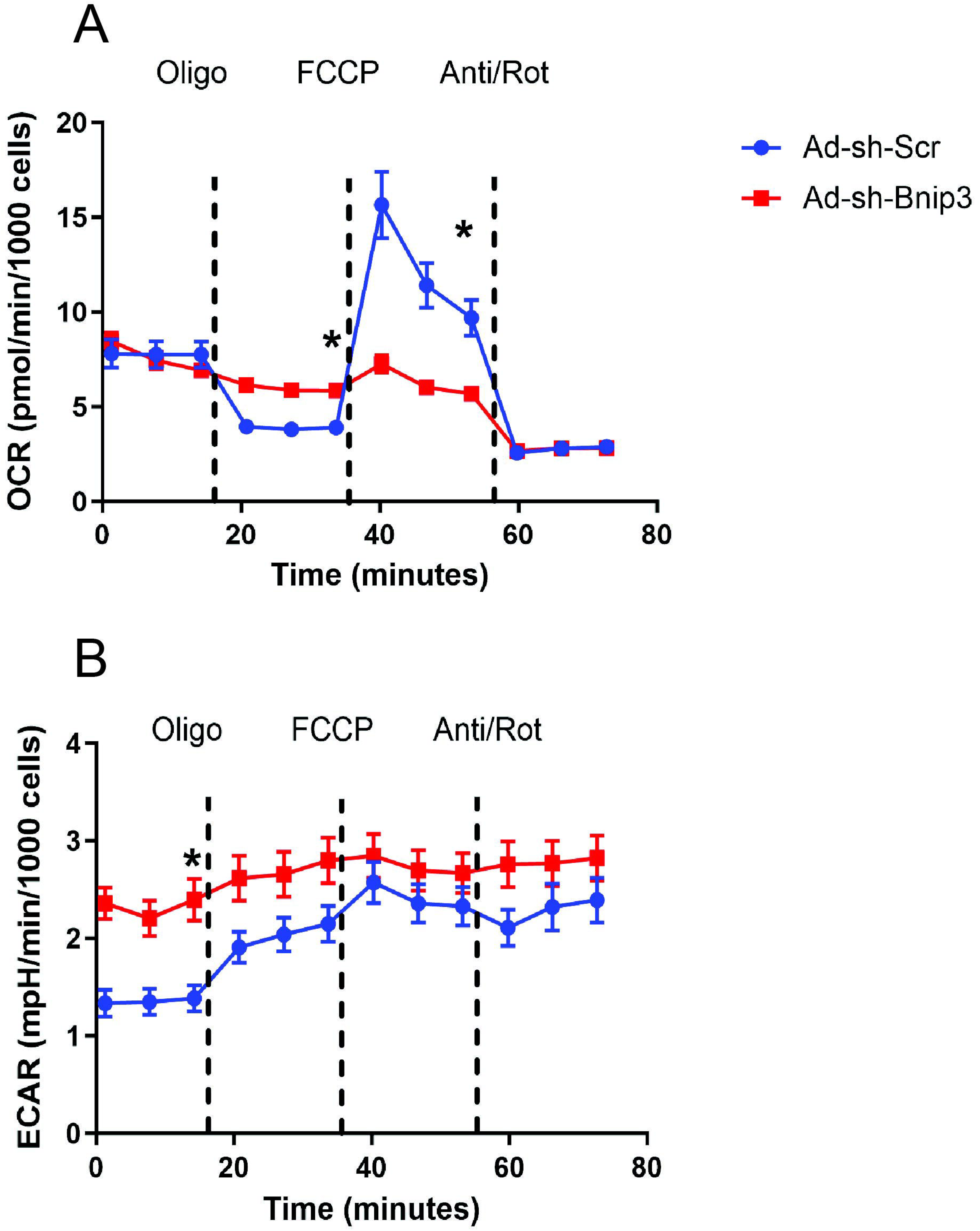
Mitochondrial dysfunction in *Bnip3* knockdown osteoblasts. **(A)** OCR (Top) and **(B)** ECAR (bottom) after Bnip3 KD (red line); blue line is scrambled control (representative of 3 trials, each with n>20 wells/group). There is increased proton leak and decreased FCCP induced OCR. There is a significant increase in basal ECAR rates as indicated in the ECAR panel before oligomycin treatment. Asterisks indicate statistically significant P-values using Student’s t-test.

### Knockout of *Bnip3* increases mitophagy and results in increased osteoblast apoptosis

Given *Bnip3’*s established role in regulating both mitophagy and apoptosis, we investigated mitophagy in the absence of *Bnip3*. To do this, we generated *Bnip3*⁻/⁻ knockout mice on a MitoQC homozygous background and isolated bone marrow stromal cells (BMSCs) from 6 to 8-week-old male mice. We then quantified mitophagy after 7days of osteogenic differentiation. Cells from the *Bnip3*⁻/⁻ group, both before and after differentiation, exhibited a significant increase in mitophagy compared to WT **(Fig 8A,B)**. Specifically, knockout cells showed a marked increase in mitophagy punctae, prompting us to examine whether they were also undergoing apoptosis. We observed an increase in p53 as shown in **(Fig 8C),** at the protein level, indicating elevated apoptosis levels. Based on these findings, we conclude that the absence of BNIP3 leads to increased mitophagy, which in turn results in heightened apoptosis in osteoblasts.

**Fig 8.**
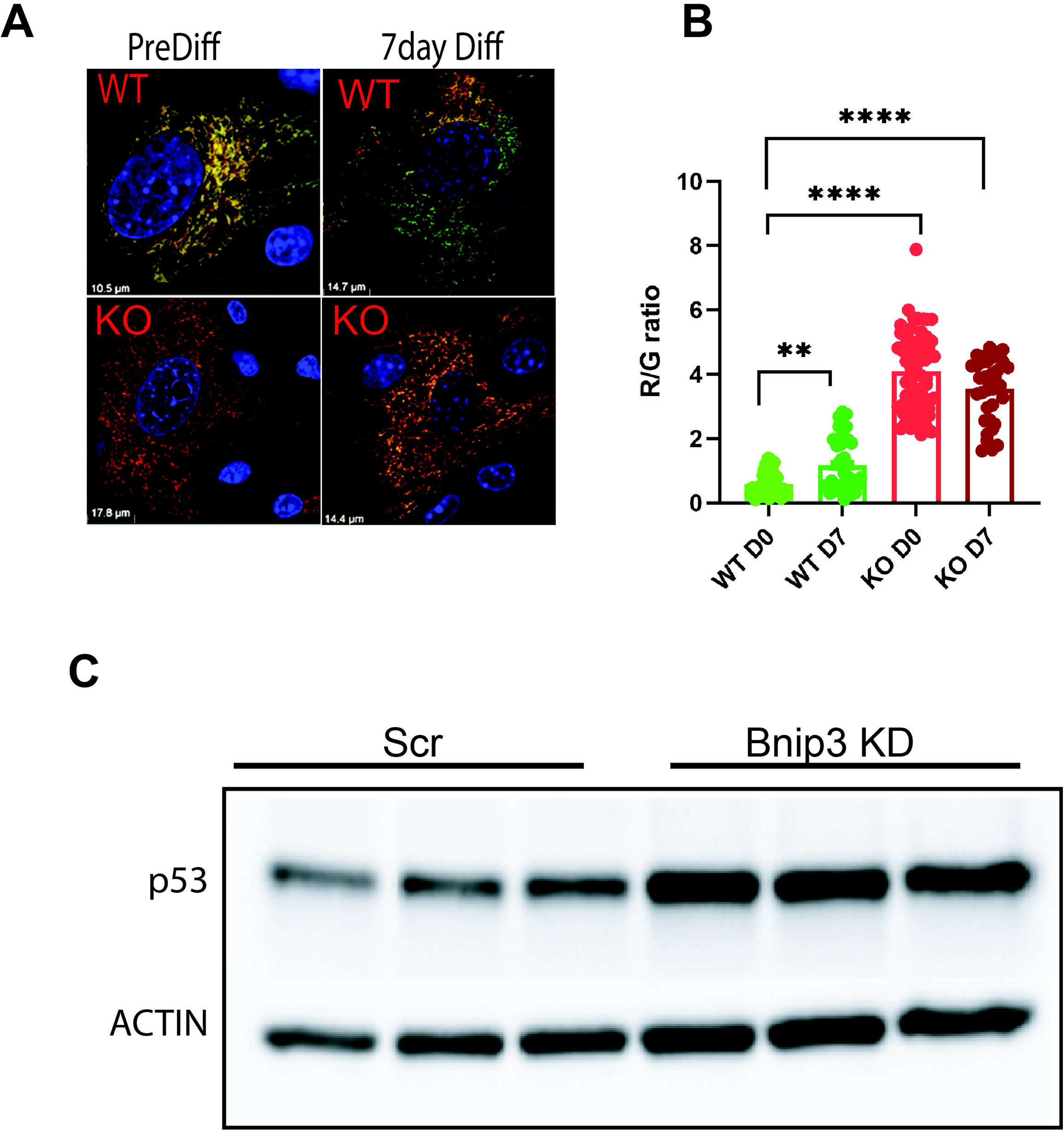
Increased mitophagy in *Bnip3*^-/-^ bone marrow stromal cells coincided with increased p53. (**A)** Mitophagy in Wildtype (WT) and *Bnip3*^-/-^ (KO) bone marrow stromal cells on a MitoQC background isolated from male mice at 6-8 wk old weeks of age. Images shown are representative of two different isolations using at least 3 animals each. BMSCs from WT and *Bnip3*^-/-^ MitoQC mice treated with non-differentiation (left panels) and 7days osteogenic differentiation (right panel). The red punctae indicate mitophagy in the top panels WT non differentiation and WT 7days and *Bnip3*^-/-^ non differentiation and *Bnip3*^-/-^ 7 days osteogenic differentiation (bottom panels). Representative confocal images from n=3 samples. **(B)** Quantification of Red/Green ratio from ≥45 cells each using MitoQC counter (FIJI), comparing WT non differentiation and WT 7days and *Bnip3*^-/-^ non differentiation and *Bnip3*^-/-^ 7 days osteogenic differentiation. P-values from one-way ANOVA followed by Tukey’s post-hoc test (\*\**p* < 0.0001). **(C)** Western blotting analysis identified an increase in p53 protein expression in *Bnip3* KD cells. ACTIN was used as the loading control.

### Mechanism of *Bnip3* mediated bone loss is through decreased ATF4 and ATF5

To identify the mechanism through which *Bnip3* mediates bone loss, we performed bulk RNA sequencing (RNA-seq) to study global gene expression changes in Adeno-scrambled control and *Bnip3* KD MC3T3E1C4 cells. Our RNA-seq data confirmed a fourfold decrease in *Bnip3* expression in our cells which were further utilized for our analysis. PCA analysis showed distinct clusters separating out scrambled controls vs *Bnip3* KD cells **(Fig 9A)**. We identified a total of 273 differentially expressed genes that are significantly modified (±1.5 fold change p≤0.05) comparing scrambled vs *Bnip3*KD cells. Further STRING analysis of the differentially expressed genes showed revealed a cluster of downregulation of multiple genes that are known targets of the bone-specific transcription factor ATF4 in our KD samples **(Fig 9B)**. These genes are also related to the integrated stress response (ISR) in relation to defects in amino acid metabolism **(Fig 9B,C)**. Cellular resilience under stress is maintained by reinitiating translation at upstream open reading frame (uORFs) in genes such as *Atf4*, *Atf5*, and *Ddit4*. Utilizing western blotting we confirmed decreases in ATF4 and ATF5 protein levels in the *Bnip3* KD cells **(Fig 9E)**. To further investigate how *Bnip3* KD affects stress responses, we treated MCT3T3E1 cells with inhibitors targeting different cellular stress pathways, Rotenone (Complex I inhibitor), Antimycin (Complex III inhibitor), Tunicamycin (Endoplasmic reticulum stress inducer) and Bafilomycin (Lysosomal acidification inhibitor). We observed a significant effect on OCR after Rotenone and Antimycin treatment, whereas OCR was relatively normal in cells treated with Tunicamycin and Bafilomycin, endoplasmic and lysosomal stress inducers respectively **(Fig 9F,G)**. This dysfunction in mitochondrial respiration was also reflected in ATF4 protein levels which were moderately upregulated in scrambled control cells and KD cells after various inhibitor treatments except for Rotenone and Antimycin which show a marked decrease in ATF4 expression, as shown in **(Fig. 9H)**. This suggests that *Bnip3* plays a role in transducing mitochondrial stress signals in response to Complex I (Rotenone) and complex III (Antimycin) inhibition to the nucleus.

**Fig 9.**
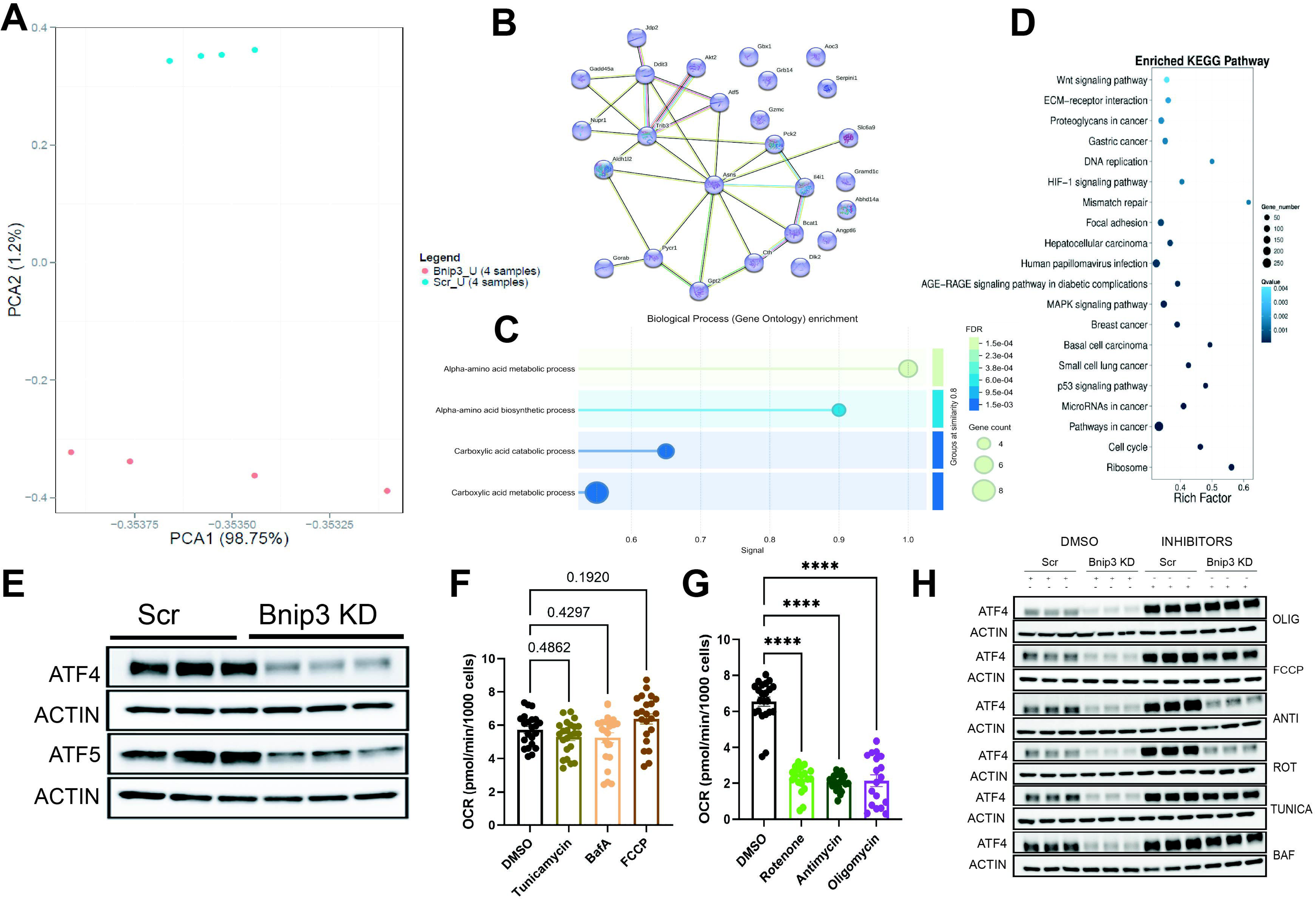
*Bnip3* knockdown induces broad transcriptional reprogramming and enrichment of stress, metabolic, and signaling pathways. **(A)** Principal component analysis (PCA) of global gene expression profiles from *Bnip3* KD (*Bnip3*, n=4) and scrambled control (Scr, n=4) MC3T3E1C4 samples. PC1 accounts for 98.75% of the total variance and clearly separates BNIP3-deficient cells from controls, indicating a strong and consistent genotype-dependent transcriptional signature. **(B)** Protein–protein interaction network of significantly altered genes generated using STRING analysis. Several key regulatory nodes, including *Atf5*, *Gadd45Aa*, *Asns*, *Akt2*, *Pycr1*, and *Slc6a9*, form a highly interconnected network associated with mitochondrial stress responses, amino acid metabolism, cell-cycle regulation, and survival pathways. **(C)** Gene Ontology enrichment (Biological Process) of BNIP3 regulated genes. Top enriched categories include regulation of metabolic processes including α-amino acid metabolic processes, α-amino acid biosynthetic process, Carboxylic acid metabolic and catabolic processes. Circle size represents gene count and color indicates false discovery rate (FDR). **(D)** KEGG pathway enrichment analysis of differentially expressed genes. Significantly enriched pathways include Wnt signaling, HIF-1 signaling, p53 signaling, MAPK signaling, ECM receptor interaction, focal adhesion, cell cycle, DNA replication, and cancer-associated pathways. The size of each bubble represents the number of genes involved and color intensity corresponds to statistical significance (q-value). **(E)** BNIP3 KD blunts ATF4 and ATF5 activation. Western blot analysis of MC3T3E1C4 cells with *Bnip3* KD compared to Scr controls showed a significant decrease in ATF4 and ATF5 levels. ACTIN was used as a loading control. **(F)** Oxygen consumption rates measured by XF96 analyzer from cells treated with DMSO, tunicamycin (2μM), bafilomycin (25nM) and FCCP (0.5µM). Asterisks indicate P-values from student’s t-test. **(G)** Oxygen consumption rates measured by XF96 analyzer show a decrease when cells were treated with rotenone (Complex I) (12μM), antimycin (Complex III) (7.6μM) and oligomycin (Complex V) (1µM). Asterisks indicate P-values from student’s t-test were significantly different between groups. **(H)** Scrambled and *Bnip3* KD cells treated overnight with rotenone (12μM) and antimycin (7.6μM), result in a stress response with an increase in ATF4 in control cells but not when BNIP3 levels are reduced, suggesting that stress responses in response to mitochondrial dysfunction require BNIP3 to transduce this signal to ATF4. Scrambled and *Bnip3* KD cells treated overnight with bafilomycin, tunicamycin, FCCP and Oligomycin which can trigger ER and lysosomal specific and mitochondrial stress response with an increase in ATF4. This response is relatively normal in the *Bnip3* KD cells with some effects from Bafilomycin and Tunicamycin suggesting that the dysfunction could partially be related to other organellar stress response. Actin is shown as loading control for each condition.

## Discussion

Osteoblast bioenergetics and substrate utilization during differentiation have been intensely studied, with conflicting results in literature. Some studies suggest that mesenchymal stem cells (MSCs) undergoing osteogenic differentiation increase reliance on oxidative phosphorylation (OxPhos), whereas others, indicate that differentiated osteoblasts rely more on aerobic glycolysis to meet the increased energetic and biosynthetic demands of matrix production^7–9^. To address mechanistic gaps underlying regulation of osteoblast bioenergetics, we first used metabolomics across multiple time points during osteogenic differentiation in MC3T3E1C4 cells.as These data revealed a coordinated, time-dependent reprogramming of cellular metabolism as differentiated cells progressively diverged from non-differentiated controls with altered glycolytic and TCA cycle intermediates, including accumulation of 2-oxoglutarate with shifts in NAD⁺/NADH linked redox balance, and evidence of disrupted mitochondrial oxidative metabolism and carbon flux. Broader metabolic remodeling was also apparent, including changes in nucleotide and amino acid metabolism and glutathione depletion indicative of increasing oxidative stress. Other published data utilizing different sources of osteogenic cells show a similar effect of differentiation on osteoblast metabolism and bioenergetics. Together, these results indicate that osteoblast differentiation imposes increasing bioenergetic, redox, and biosynthetic demands that are met through broad rewiring of central metabolism and nucleotide homeostasis^19^.

Based on these observations, and supported by recent work showing that differentiating cells can upregulate mitophagy in a programmed manner^20^, we next examined mitophagy using the MitoQC transgenic reporter model. We identified basal mitophagy in skeletal cells in vivo and in isolated calvarial osteoblasts, and observed that mitophagy increased as osteogenic differentiation progressed. Importantly, increased mitophagy coincided with increased expression of the mitophagy receptor *Bnip3*, with a smaller increase in *Nix/Bnip3l*. Although correlative links between mitochondrial turnover and skeletal remodeling have been suggested, mechanistic data connecting mitochondrial quality control to osteoblast differentiation and mineralization remains limited. Our findings place receptor-associated mitophagy, and BNIP3 specifically, as a key node linking differentiation-associated metabolic remodeling to mitochondrial quality control in osteoblasts.

BNIP3 is a ∼28 kDa protein that localizes to the outer mitochondrial membrane via its C-terminus and, through an N-terminal LC3-interacting region (LIR), can function as a mitophagy receptor. BNIP3 can form homodimers, interact with proteins such as BCL2^21^. BNIP3 was initially described as a pro-apoptotic factor through a BH3-like domain, but more recent work has established its role as a mitophagy receptor that binds LC3II and recruits adaptor proteins required for initiating mitochondrial autophagy^18^. BNIP3 and NIX dysregulation in humans is associated with mitochondrial DNA depletion syndrome (MTDPS13), where stabilization of BNIP3 and NIX results in increased mitochondrial turnover and dysfunction^5–7^, leading to encephalomyopathy, short stature and developmental defects. Disease etiology includes mutations in the E3 ubiquitin ligase complex, SCF-FBLX4 which normally targets BNIP3 and NIX to the proteasome for degradation. When FBLX4 is mutated or depleted in MTDPS13, BNIP3 and NIX are stabilized, resulting in increased mitochondrial turnover and dysfunction. We have previously shown that BNIP3 is downstream of HIF1α signaling in intervertebral discs and that loss of BNIP3 affects mitophagy and disc anatomy through regulation of cellular metabolism. In bone, prior work in C57BL/6J mice reported decreased *Bnip3* expression with aging^22^. Together, these observations support the idea that BNIP3 is positioned to coordinate mitochondrial homeostasis with metabolic state and stress signaling in differentiated tissues, including skeletal cells^23^.

To test the functional requirement for BNIP3, we used adenoviral shRNA-mediated KD and found reduced expression of osteoblast differentiation genes in vitro, with decreased mineralization. Proteomic analyses further demonstrated broad disruption of proteins and pathways related to mitochondrial organization, protein localization to organelles, translation, and cellular metabolism, collectively indicating that BNIP3 loss disrupts mitochondrial organization, metabolic homeostasis, and biosynthetic regulation necessary for osteogenic maturation. Consistent with these molecular signatures, BNIP3 loss was associated with defective mitochondrial oxidative phosphorylation characterized by reduced spare respiratory capacity, increased proton leak, and a compensatory shift toward glycolysis, consistent with mitochondrial uncoupling and bioenergetic stress.

In vivo, *Bnip3* global knockout animals have been reported to have metabolic defects in liver and adipose tissue that could theoretically influence bone through inter-organ crosstalk^24^. We note this potential endocrine contribution and address it as a future direction using a conditional *Bnip3* allele. Despite potential systemic effects, we observed a clear bone phenotype in *Bnip3⁻/⁻* male mice. µCT analysis revealed significant decreases in trabecular bone mass and cortical expansion with reduced polar moment of inertia, indicating weaker bones. The phenotype was sexually dimorphic, with females showing minimal skeletal changes compared to controls. Histomorphometry in males supported a cellular basis for low bone mass reduced osteoblast number and reduced measures of bone formation, without changes in osteoclast number, marrow adiposity, or systemic bone turnover markers. These results indicate that low bone mass in *Bnip3⁻/⁻* mice arises predominantly from impaired osteoblast number and activity rather than increased bone resorption. The sexually dimorphic phenotype suggests that females may be protected by estrogen and/or compensatory mechanisms, potentially involving other mitophagy factors such as *Bnip3l* and *Bcl2l13*, which increased during osteogenesis.

A central finding from our study is that BNIP3 links mitochondrial dysfunction to nuclear stress-response programs in osteoblasts. Mechanistically, we identified a defect in the classical integrated stress response and the mitochondrial stress response mediators ATF4 and ATF5 in BNIP3 deficient cells. Mitochondrial dysfunction is a major causal factor in aging-related disorders, including osteoporosis, and mitochondria are recycled through mitophagy, which includes the PINK/PARKIN ubiquitination pathway, receptor-mediated mitophagy, and mitochondrial-derived vesicles^3^. Defects in mitophagy are implicated in multiple human diseases that are associated with skeletal fragility, including lysosomal storage diseases and Barth syndrome, and skeletal defects are also associated with human mitochondrial DNA mutations^25,26^. In relevant osteoblast settings, mitochondrial stress responses can be engaged independent of classical ER stress, as shown in models where ATF5 increases to compensate for procollagen misfolding^27^. These observations support the concept that osteoblasts utilize mitochondria-linked stress-response pathways to preserve function under biosynthetic load necessary for driving its major cellular function of collagen secretion and mineralization.

In our system, ATF4 and ATF5 were downregulated in *Bnip3* KD cells. ATF4 is critical for skeletal development and, beyond its role in osteoblast function, is a canonical effector of the integrated stress response (ISR)^28–32^. ATF4 is selectively translated downstream of eIF2α phosphorylation by stress-responsive eIF2 kinases, including GCN2, which is activated by amino acid deficiency and regulates osteoblast progenitor proliferation upstream of ATF4 to control glutamine metabolism^33, 34^. ATF4 can also increase import of amino acids relevant to collagen synthesis^47^.

In our study, while we did not observe a strong change in pSer52-eIF2 status, our transcriptomic and proteomic data indicated defects in amino acid metabolism and protein synthesis programs. RNA-seq and STRING-based network analysis identified a cluster of ATF4-regulated genes that were downregulated in *Bnip3* KD cells, accompanied by decreased ATF4 and ATF5 at the protein level. We also observed decreased expression of multiple cytosolic aminoacyl tRNA synthetases, consistent with reduced amino acid metabolism and translational capacity. Because collagen synthesis is a major function of osteoblasts, these defects provide a plausible mechanism by which impaired stress signaling and biosynthetic support contribute to reduced osteoblast function.

To probe whether the ATF4 defect reflected a mitochondria specific signaling impairment, we challenged cells with perturbations targeting mitochondrial electron chain complexes (Complex I and Complex III) and other organellar stress pathways. We observed marked dysfunction in OCR following rotenone (Complex I inhibitor) and antimycin (Complex III inhibitor) treatments, whereas responses to tunicamycin (ER stress) and bafilomycin (lysosomal stress) were comparatively preserved. Correspondingly, ATF4 was moderately induced in scrambled control cells after several inhibitor treatments, but rotenone and antimycin were associated with a marked decrease in ATF4 when BNIP3 levels were reduced. These observations support a model in which BNIP3 is required to transduce mitochondrial stress signals to ATF4/ATF5-dependent nuclear programs, while responses to non-mitochondrial stressors are relatively less affected.

Finally, BNIP3 deficiency was associated with increased mitophagy and increased activation of p53. Together with the in vivo finding of reduced osteoblast number and reduced bone formation, these data support a model where during osteoblast differentiation, BNIP3 is induced alongside increased mitophagy, and BNIP3 is necessary not only for mitochondrial homeostasis and oxidative capacity but also for mitochondrial stress to nuclear signaling through ATF4 and ATF5. Loss of this BNIP3 dependent relay impairs the ability of osteoblast lineage cells to mount appropriate mitochondrial stress responses and maintain biosynthetic and survival programs, resulting in increased apoptosis, reduced osteoblast numbers and activity, and decreased bone mass in male mice.

### Future directions and limitations

Future work will delineate the role of *Bnip3*, specifically in osteoblast and osteocyte function. The increase seen in other mitophagy receptors *Bcl2l13* and *Bnip3L* at the gene expression level during differentiation suggest some potential role for these factors in bone. We have previously studied *Bcl2l13* in adipogenic differentiation and its role in regulating mitophagy during adipogenesis suggesting that here are still other potential factors that need to be identified that can regulate mitophagy in osteoblasts. We are currently pursuing changes we observed in metabolites during osteogenic differentiation and if they have any functional consequences on osteoblasts through epigenetic modifications.

## Materials and methods

### Animals

MitoQC transgenic mice were obtained from Dr. Ian Ganley at the University of Dundee and we have published previously with these mice ^35^. *Bnip3^-/-^* animals were generously provided by Dr.Gerald Dorn II and were described previously^24^. These mice were rederived at Washington University Medical Center at St. Louis (WASHU) and shipped to MHIR. All animals used were housed at MaineHealth Institute for Research (MHIR) mouse vivarium, a germ free barrier facility on a 14hr light and 10hr dark schedule. Animals had free access to water (autoclaved) and food (Teklad global 18% protein diet, #2918, Envigo, breeder diet). All animal procedures described in this study were approved by the Institution Animal Care and Use Committee (IACUC) at MHIR.

### Cell culture

#### MC3T3E1 culture

MC3T3E1 sub clone 4 (MC3T3E1C4) was obtained from (ATCC) and cultured in α-MEM supplemented with 10% fetal bovine serum, 100U/ml penicillin and 100mg/ml streptomycin. Differentiation was initiated by switching culture media to osteoblast (OB) differentiation medium containing 50µg/ml ascorbic acid and 8mM β-glycerophosphate. OB differentiation medium was changed every other day.

#### Primary calvarial osteoblast (COB) isolation and culture

Neonate calvarial osteoblasts were isolated from 1-3-day *Bnip3*^-/-^ mice and control (WT) littermate mice. In brief, the calvarial osteoblasts were subjected to four sequential digestions with Collagenase P (1.6U/mg) and 0.25% trypsin/EDTA on shaker at 37^0^C for 15 min. Digests 2-4 were collected and plated as described previously^6^.

#### Bone marrow stromal cell (BMSCs) isolation and culture

Hindlimbs (both femurs and tibiae) from 6-8 week old male or female mice were dissected and bone marrow was spun out into micro centrifuge tubes (∼13000rpm for 15seconds at RT) as described previously^18^. BMSC cultures were plated and maintained in α-MEM supplemented with 10% fetal bovine serum, 100U/ml penicillin and 100mg/ml streptomycin. Differentiation was induced with osteoblast (OB) differentiation medium containing 50µg/ml ascorbic acid and 8mM β-glycerophosphate. OB differentiation medium was changed every other day.

### Confocal microscopy and mitophagy quantification

To observe mitophagy in primary calvarial osteoblast from MitoQC mice, we plated the primary calvarial osteoblasts with a density of 1×10^5^/well on cover slips. After cells reached confluence defined as 0d, cells were washed with 1X PBS and fixed with 4% paraformaldehyde for 10 min. Fixed cells were processed for mounting using Vector DAPI fluorescence mounting medium and a Leica SP8 confocal microscope was utilized to capture images. Osteoblast differentiation was followed as described above for 7d, 14d, and 21d. Quantification was performed on images captured using a 63x oil immersion objective and processing data using Mito counter as described in^36^ and mitophagy is represented as Red to green ratio.

### Adenoviral shRNAi knockdown

Adenovirus mediated short hairpin RNA (shRNAi) against mouse *Bnip3* (Ad-sh-*Bnip3*) and a negative control scrambled shRNAi (Ad-sh-Scr) were purchased from Vector Biolabs (PA, USA). MC3T3E1C4 cells were plated in a 6-well tissue culture plate at a density of 1.2×10^5^/well. Cells were cultured for at least 15h in α-MEM supplemented with 10% fetal bovine serum, 100U/ml penicillin and 100mg/ml streptomycin prior to virus transfection. The cells were infected when they reached 50-60% confluence at a multiplicity of infection (MOI) of 100 in serum free, antibiotic-free α-MEM medium. After 12h infection at 37^0^C media was exchanged to fresh complete medium. KD was confirmed at both the transcript and protein level.

### Chemicals

Deferiprone (Sigma#379409) treatment was done overnight at a concentration of 1mM in methanol; CoCl2 (Sigma#158621) was used at a concentration of 200µM, Bafilomycin (Sigma#1661) was used at a concentration of 25nM, Mitotracker Red (Cell Signaling Technology #9082) was used at a concentration of 100nM for 45 min at 37^0^C before fixation and confocal imaging as described in the section above.

### Agilent XF96 Seahorse assays

MC3T3E1C4 cells were seeded in 96-well Seahorse XF cell culture microplate at a density of 2.5×10^3^/well in complete medium. Cell were cultured and infected at MOI=100 as described above. 12h post infection cells were switched to complete growth media for 72h and this timepoint was defined as undifferentiated condition. For differentiation, we switched cells to OB differentiation medium every other day until the 7d, or 14d. We measured oxygen consumption rate (OCR) and extracellular acidification rate (ECAR) with the Seahorse XFe Extracellular Flux Assay Kit in a Seahorse XFe96 Analyzer (Agilent Technologies, USA), following the manufacturer instruction and this protocol has been described by us previously. The concentrations of reagents used for the mitostress test have been described previously^10^.

### Gene expression analysis

Total RNA from cells and tissue was extracted using TRIzol reagent (Invitrogen, USA) following manufacturer instructions. cDNA was synthesized via reverse transcription using the High Capacity cDNA Kit (Applied Biosystems). The relative gene expression level was quantified with iQ SYBR Green Supermix (Bio-Rad, USA) in the CFX Connect Real-Time PCR Detection System. The fold change of gene expression was analyzed by the 2^-ΔΔCt^ method and normalized to the housekeeping gene hypoxanthine phosphoribosyltransferase (HPRT) expression shown as relative expression. The primer sequence list used in this study are listed in **(Supplemental table 1)**.

### Microcomputed tomography (µCT)

Soft tissue was removed and the bones fixed in 70% ethanol. The distal end of femurs were scanned using standard μCT techniques. A high-resolution desktop micro-tomographic imaging system (µCT40, Scanco Medical AG, Brüttisellen, Switzerland) was used to assess trabecular bone architecture in the distal femoral metaphysis and cortical bone morphology of the femoral mid-diaphysis. Scans were acquired using a 10 µm^3^ isotropic voxel size, 70 kVP, 114 μA, 200 ms integration time, and were subjected to Gaussian filtration and segmentation. Image acquisition and analysis protocols adhered to the JBMR guidelines for the assessment of rodent bones using µCT. In the femur, trabecular bone microarchitecture was evaluated in a region beginning 1.3% of the total femur length superior to the distal growth plate and extending proximally 10% of the femur length. The lengths of the trabecular regions of interest ranged from 420 µm for the shortest femur to 1,570 µm for the longest femur. The trabecular bone regions were identified by manually contouring the endocortical region of the bone. A threshold of 375 mgHA/cm^3^ was used to segment bone from soft tissue. Trabecular microarchitecture was analyzed using the standard trabecular bone morphology script in the Scanco Evaluation Program. The following architectural parameters were measured using the Scanco Trabecular Bone Morphometry evaluation script: trabecular bone volume fraction (Tb.BV/TV, %), trabecular bone mineral density (Tb. BMD, mgHA/cm^3^), specific bone surface (BS/BV, mm^2^/mm^3^), trabecular thickness (Tb.Th, mm), trabecular number (Tb.N, mm^-1^), and trabecular separation (Tb.Sp, mm), connectivity density (Conn.D, 1/mm^3^), and structural model index (SMI. Cortical bone was assessed in 50 transverse µCT slices (500 μm long region) at the femoral mid-diaphysis and the region included the entire outer most edge of the cortex. Cortical bone was segmented using a fixed threshold of 700 mgHA/cm^3^. The following variables were computed: total cross-sectional area (bone + medullary area) (Tt.Ar, mm^2^), cortical bone area (Ct.Ar, mm^2^), medullary area (Ma.Ar, mm^2^), bone area fraction (Ct.Ar/Tt.Ar, %), cortical tissue mineral density (Ct.TMD, mgHA/cm^3^), cortical thickness (Ct.Th, mm), as well as maximum, minimum and polar moments of inertia (Imax, Imin, and J, mm^4^), which describe the shape/distribution of cortical bone (larger values indicate a higher bending strength).

### Bone histomorphometry

Bone histomorphometric analysis was performed on femurs from 16 weeks of age and sex-matched controls. In brief, calcein (10mg/kg) and alizarin (30mg/kg) were injected at 7 days and 2 days prior to animal euthanasia, respectively. Samples were dissected, cleaned free of soft tissue, immersed in 70% ethanol, and processed as previously described^20^. Non-decalcified femurs were dehydrated (graded ethanol) and subsequently infiltrated and embedded in methylmethacrylate. Longitudinal sections (5 micron) were cut using a microtome (RM2255, Leica) and stained with Goldner’s Trichrome for measurements of cellular parameters. Dynamic bone parameters were evaluated on unstained sections by measuring the extent and the distance between double labels using a semi-automatic image analyzer system (Bioquant Osteo 2020). Measurements were made in the area 200-250 mm proximal to the distal growth plate and at least two nonconsecutive sections per sample were examined. The structural, dynamic, and cellular parameters were evaluated using standardized guidelines^37^.

### Western blotting

Protein lysates were prepared in RIPA buffer (Cell Signaling Technology,Inc., Danvers, MA, USA#) containing protease inhibitor cocktail (Roche, Basel, Switzerland#). Protein concentrations were determined using BCA (Pierce#) commercially available assay kit : 40µg protein were loaded on the 4-15% TRIS-Glycine Mini Gels (Bio-Rad), and run at 80 V for 30 minutes, followed by 130V for 50 minutes, and then transferred to a PVDF membrane. The membrane was blocked in 5% non-fat milk in TBST (Tween 20 0.1%). Primary and secondary antibodies were incubated in 5% non-fat milk in TRIS buffered saline with Tween 20 (TBST) buffer. Protein bands were visualized using SuperSignal West Pico Chemiluminescence substrate (Thermo Fisher# 34580) and imaged with a ChemiDoc Touch Imaging System (Bio-Rad). Scanned images were analyzed with Image J and the protein relative expression was normalized to β-actin or Cyclophilin A. Experiments were repeated at least 3 times. The following antibodies were used: polyclonal rabbit anti-BNIP3 (1:1000; #3769, Cell Signaling Technology), monoclonal mouse anti-beta-Actin(C4) (1:1000; # sc-47778, Santa Cruz Biotechnology). ATF4 (1:1000; #11815, Cell Signaling Technology), ATF5 (1:1000; #3769, Cell Signaling Technology), P53(1:1000; #9282, Cell Signaling Technology).

### Cell Mineralization assay

The protocol utilized has been described in ^6^. Cell mineralization was assessed by OsteoImage mineralization fluorescent assay (Lonza, Switzerland# PA-1503) following the manufacturers protocol. For the OsteoImage mineralization fluorescent assay readings were obtained on a cyation instrument.

### Bulk RNA sequencing analysis BGI workflow

Briefly mRNA was isolated from 4 scrambled control and 4 *Bnip3* KD samples each (using oligo(dT)-attached magnetic beads, followed by fragmentation of the mRNA into smaller pieces under controlled conditions. First-strand cDNA is synthesized using random hexamer primers, and second-strand cDNA synthesis follows. The cDNA fragments are then end-repaired, 3’ adenylated, and ligated with sequencing adapters. These adapter-ligated fragments are amplified via PCR, purified using Ampure XP beads, and dissolved in EB solution. The resulting libraries are assessed for quality using the Agilent 2100 Bioanalyzer. Subsequently, double-stranded PCR products are heat-denatured and circularized with splint oligonucleotides, forming single-strand circular DNA (ssCir DNA), which serves as the final sequencing library. Amplification of these libraries using phi29 polymerase generates DNA nanoballs (DNBs), each containing more than 300 copies of the original molecule. These DNBs were loaded onto a patterned nanoarray, and sequencing was performed using combinatorial Probe-Anchor Synthesis (cPAS), producing single-end 50 bp or paired-end 100/150 bp reads. Data analysis was performed by BGI.

### Proteomics

The proteins immobilized on the NxStage filters were analyzed by mass spectrometry in the laboratory of Dr. Simone Sidoli at Albert Einstein College of Medicine (AECM). Briefly, protein lysates were frozen at MHIR and shipped to Dr. Sidolis laboratory. Protein lysates were then extracted using organic solvents, dried in a SpeedVac and digested using trypsin. Proteomics identification and quantification was performed using state-of-the-art mass spectrometry (Orbitrap Fusion Lumos, Thermo Scientific). Spectra processing, peptide identification, peptide quantification, and protein roll-up (i.e., quantification of proteins using peptide intensities) was processed by Proteome Discoverer (v2.4, Thermo Scientific). Data transformation, normalization, and statistics were performed using BigOmics software analysis to do PCA, analysis pathway analysis and enrichment analysis. Protein lysates were prepared from n=3 samples each (scrambled control vs *Bnip3* KD).

### Metabolomics

Cultured cells: To investigate intracellular metabolite dynamics during osteoblast differentiation, we utilized MC3T3E1C4 pre-osteoblast clones and performed metabolomic studies. Cells were cultured in growth media and osteogenic differentiation media for 7, 14, and 19 days. Intracellular metabolites were extracted using the protocol described previously^38^. Additionally, metabolite extraction followed a standardized protocol, involving rapid quenching with pre-chilled 80% methanol solution cell scraping, centrifugation, and collection of the metabolite-containing supernatant for downstream analysis. Metabolites were analyzed using an unlabeled polar metabolite profiling method on a 5500 QTRAP hybrid triple quadrupole mass spectrometer, covering major metabolic pathways at the core facility at BIDMC. Cell culture metabolomics data were processed using Gigantest’s Laboratory Information Management System (LIMS) in combination with Python, R and JavaScript. Specifically, first, fold change was calculated by dividing the intensity of each metabolite for each sample by the average intensity of the control group for that metabolite. p-values were calculated using a one-tailed student’s t-test, for every pairwise comparison, Volcano plots were plotted between each pair of experimental groups, using metabolite intensities, integrating both fold change and statistical significance (with a p-value threshold of 0.05). Metabolites which are higher and statistically significant are in red dots. Metabolites which are lower and statistically significant are in blue dots. Metabolites which are not statistically significant are in black dots. The positioning of each dot in the plot is indicative of both the magnitude of change and the statistical significance. Dots positioned farther away from the origin signify greater differences both in terms of absolute value and statistical relevance. PCA plots were plotted using metabolite intensities, in which experimental group names were blinded to assess the variance in the data and samples clustered based on the metabolic similarities within the dataset^39, 40^.

### Statistics

The data is represented as mean ± standard error of the mean (SEM). Comparison between two groups was done using the student *t* test. One-way ANOVA was used for the multiple group’s comparison. P value ≤ 0.05 is significant. All the data was analyzed using GraphPad version. Mitophagy quantification was done by using SPSS 20.0 (IBM Inc., Armonk, NY, USA).

## Acknowledgements

This work was supported by funds from NIH R01HD112474, NIH/NIGMS, P20GM121301 and funds from MaineHealth Institute for Research (MHIR) to GAR. This work utilized services of the following core facilities at MHIR, Molecular Phenotyping Core, which is supported by NIH/NIGMS P30GM106391, the Physiology Core, supported by NIH/NIGMS P30GM106391 and P20GM121301, and the Mouse Transgenic and In Vivo Imaging Core, which is supported by NIH/NIGMS P30GM103392. All cores also received support from the Northern New England Clinical and Translational Research Network NIH/NIGMS U54GM115516. We thank Dr. Gerald Dorn II (WASHU) for sharing the *Bnip3^-/-^* mice. We thank Dr. Carolyn Chlebek for help with editing the manuscript. We acknowledge using ChatGPT version 5 for better language clarity and flow during preparation of this manuscript. The content is solely the responsibility of the authors and does not necessarily represent the official views of the National Institutes of Health.

## Data availability

The data supporting the findings from this study are available from GAR upon reasonable request.

