## Supplementary Fig for "Impaired mitochondrial stress signaling mediates bone loss in male mice in the absence of BNIP3"

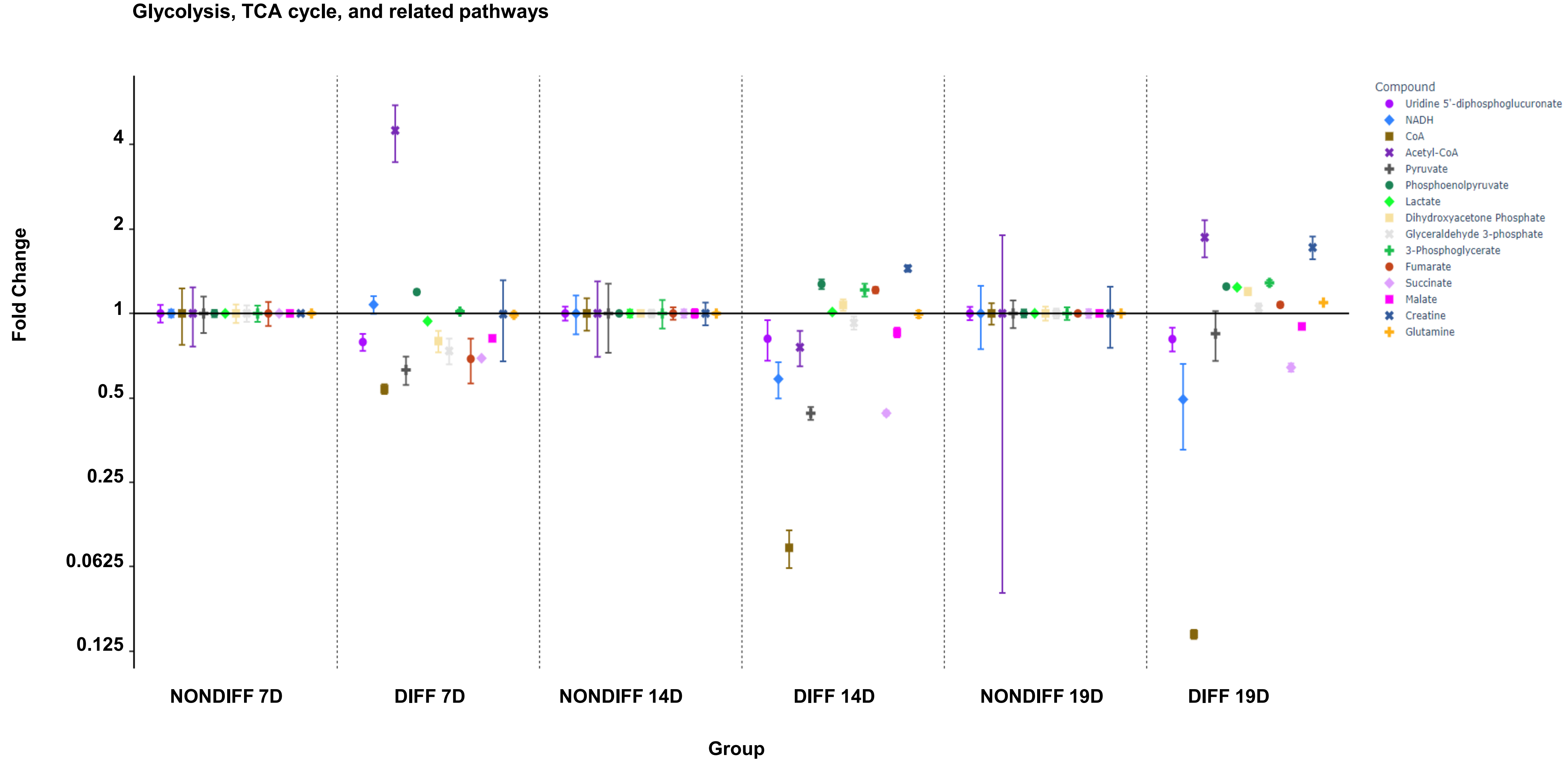


SFig1: Altered glycolytic and related metabolic flux in response to differentiation: The panel shows relative fold changes in selected metabolites associated with glycolysis and the TCA cycle across experimental groups (NONDIFF_7d, DIFF_7d, NONDIFF_14d, DIFF_14d, NONDIFF_19d, DIFF_19d). Each point represents the mean abundance of a specific metabolite with error bars indicating variability. Compounds displayed include NADH, $NAD^+$, acetyl-CoA, pyruvate, phosphoenolpyruvate, lactate, dihydroxyacetone phosphate, glyceraldehyde-3-phosphate, 3-phosphoglycerate, fumarate, succinate, malate, creatine, and glutamine. Overall, the data reveal dynamic, time-dependent shifts in central carbon metabolism during differentiation, with more pronounced deviations in differentiated samples

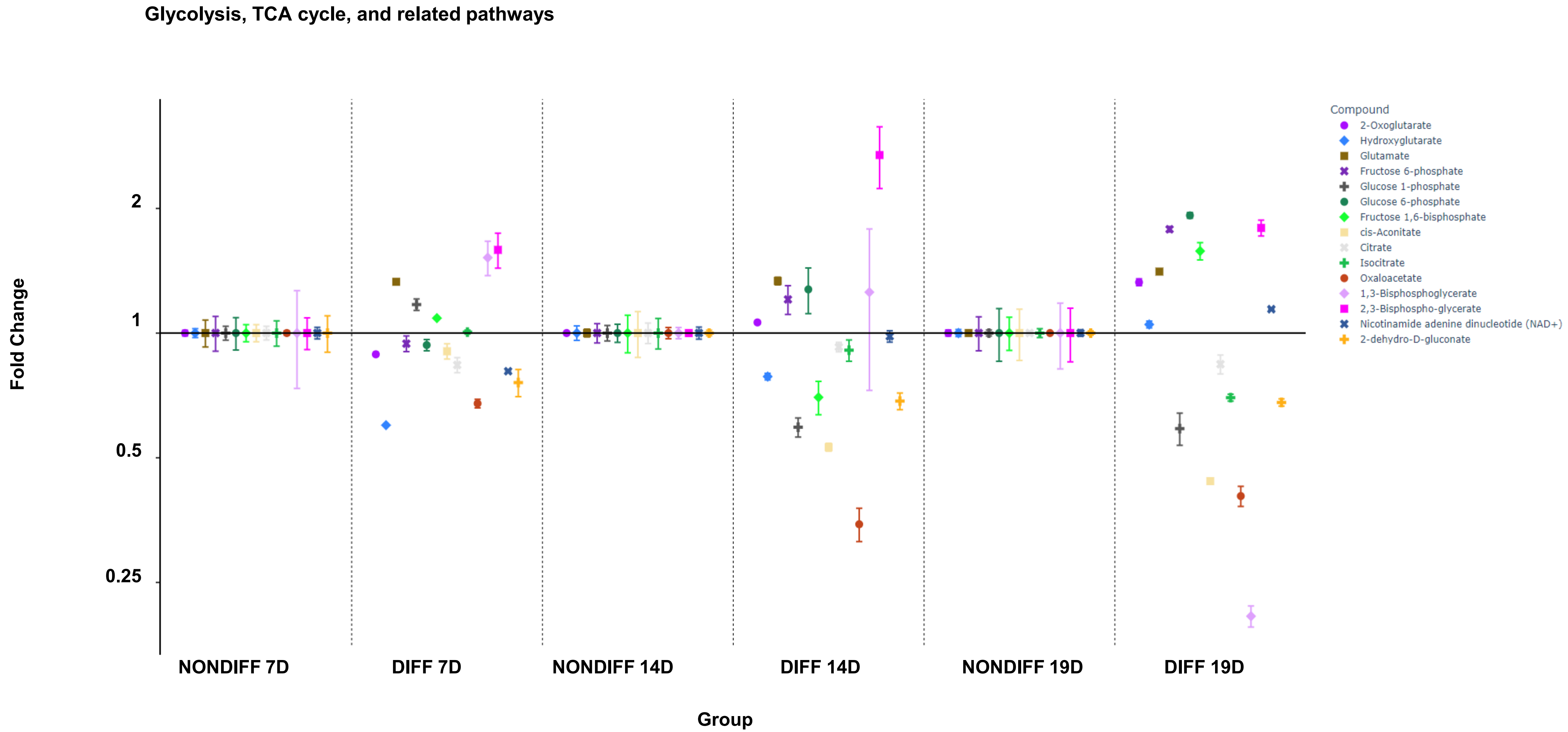


SFig2:The panel shows relative fold changes of metabolites associated with the TCA cycle and closely linked glycolytic intermediates across experimental groups (NONDIFF_7d, DIFF_7d, NONDIFF_14d, DIFF_14d, NONDIFF_19d, DIFF_19d). Each colored point represents a specific metabolite, with error bars indicating variability between replicates. Metabolites displayed include α-ketoglutarate, hydroxyglutarate, glutamate, citrate, cis-aconitate, isocitrate, oxaloacetate, 1,3-bisphosphoglycerate, 2,3-bisphosphoglycerate, nicotinamide adenine dinucleotide ($NAD^+$), and 2-dehydro-D-gluconate. Several intermediates show time- and differentiation-dependent increases or decreases, indicating altered carbon flux through the TCA cycle. The observed changes in TCA intermediates, along with shifts in $NAD^+$/NADH-related metabolites, support a disruption in mitochondrial oxidative metabolism and anaplerotic/cataplerotic balance during osteoblast differentiation.

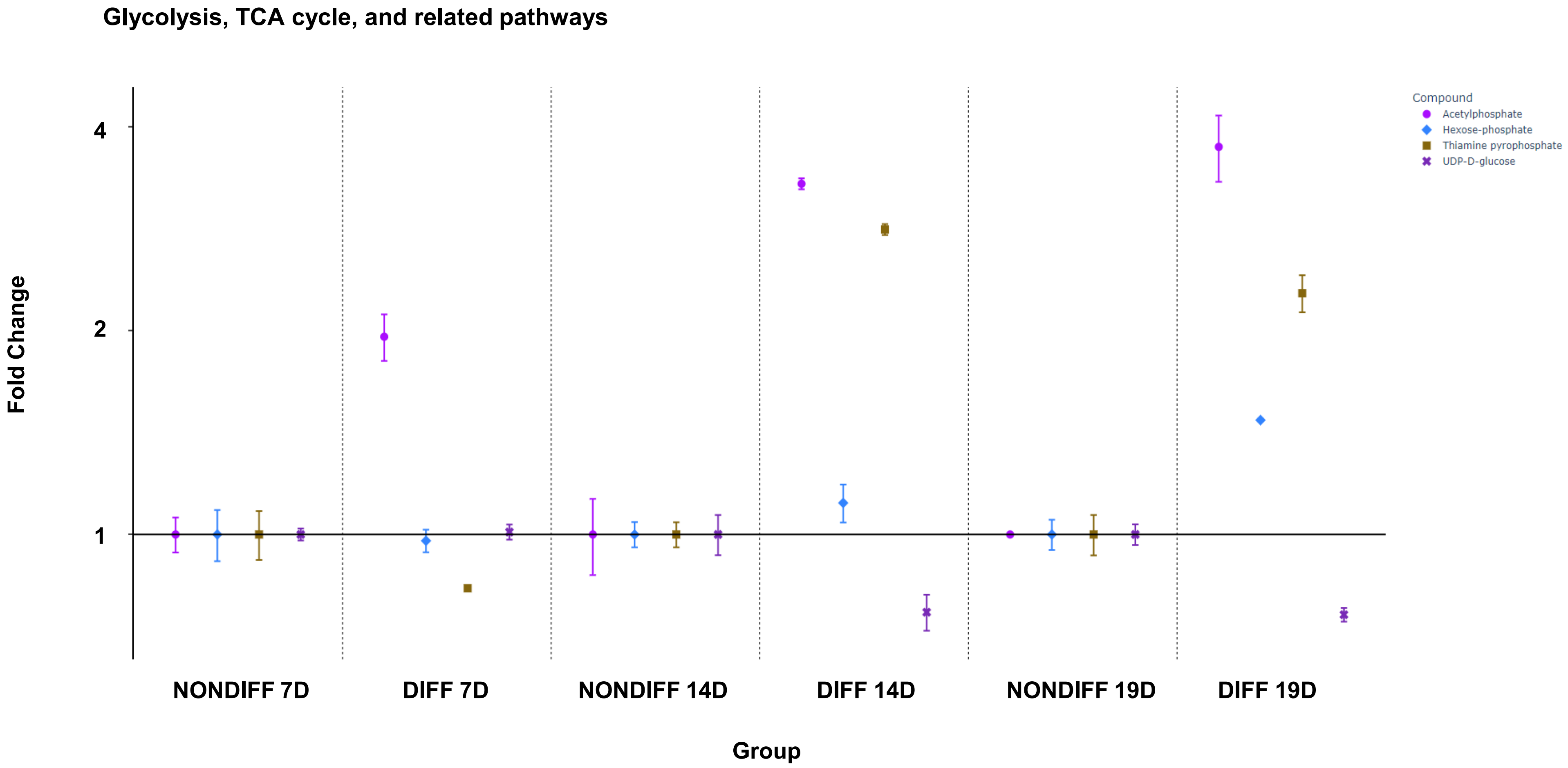


SFig3 :Relative fold changes of selected metabolites are shown across experimental groups (NONDIFF_7d, DIFF_7d, NONDIFF_14d, DIFF_14d, NONDIFF_19d, DIFF_19d). Compounds displayed include acetylphosphate, hexose-phosphate, thiamine pyrophosphate (ThPP), and UDP-D-glucose. Each point represents the mean fold change relative to baseline, with error bars indicating variability. While non-differentiated groups remain near baseline, differentiated samples exhibit marked, time-dependent deviations most prominently at 14 and 19 days indicating progressive reprogramming of central carbon metabolism. The increase in thiamine pyrophosphate and acetylphosphate, together with altered hexose-phosphate and UDP-glucose levels, suggests shifts in glycolytic flux, cofactor availability, and anabolic carbohydrate metabolism

### De-Novo Purine Synthesis

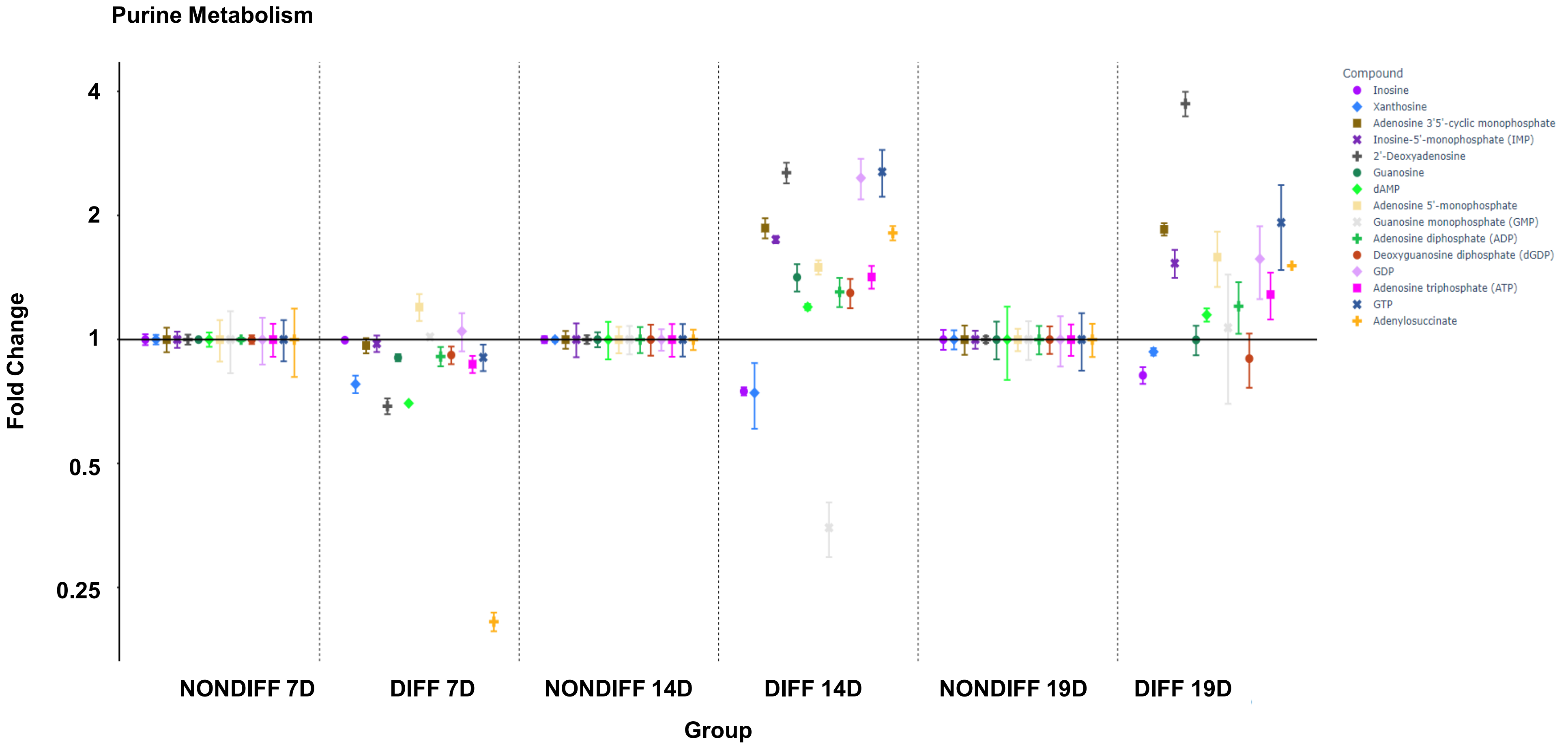


SFig4:Modulation of de novo purine synthesis during osteogenic differentiation. The panel displays relative fold changes in purine and nucleotide intermediates across experimental groups (NONDIFF_7d, DIFF_7d, NONDIFF_14d, DIFF_14d, NONDIFF_19d, DIFF_19d). Compounds include inosine, xanthosine, IMP, AMP, ADP, ATP, GMP, GDP, GTP, adenosine, deoxyadenosine, deoxyadenosine diphosphate, adenosine 3',5'-cyclic monophosphate, and adenylosuccinate. While non-differentiated cells remain near baseline, differentiated samples show time dependent accumulation of several purine intermediates and nucleotides, with particularly pronounced increases at later time points. These data suggest enhanced nucleotide demand and/or altered purine turnover in response to differentiation.

De-novo Pyrimidine Synthesis

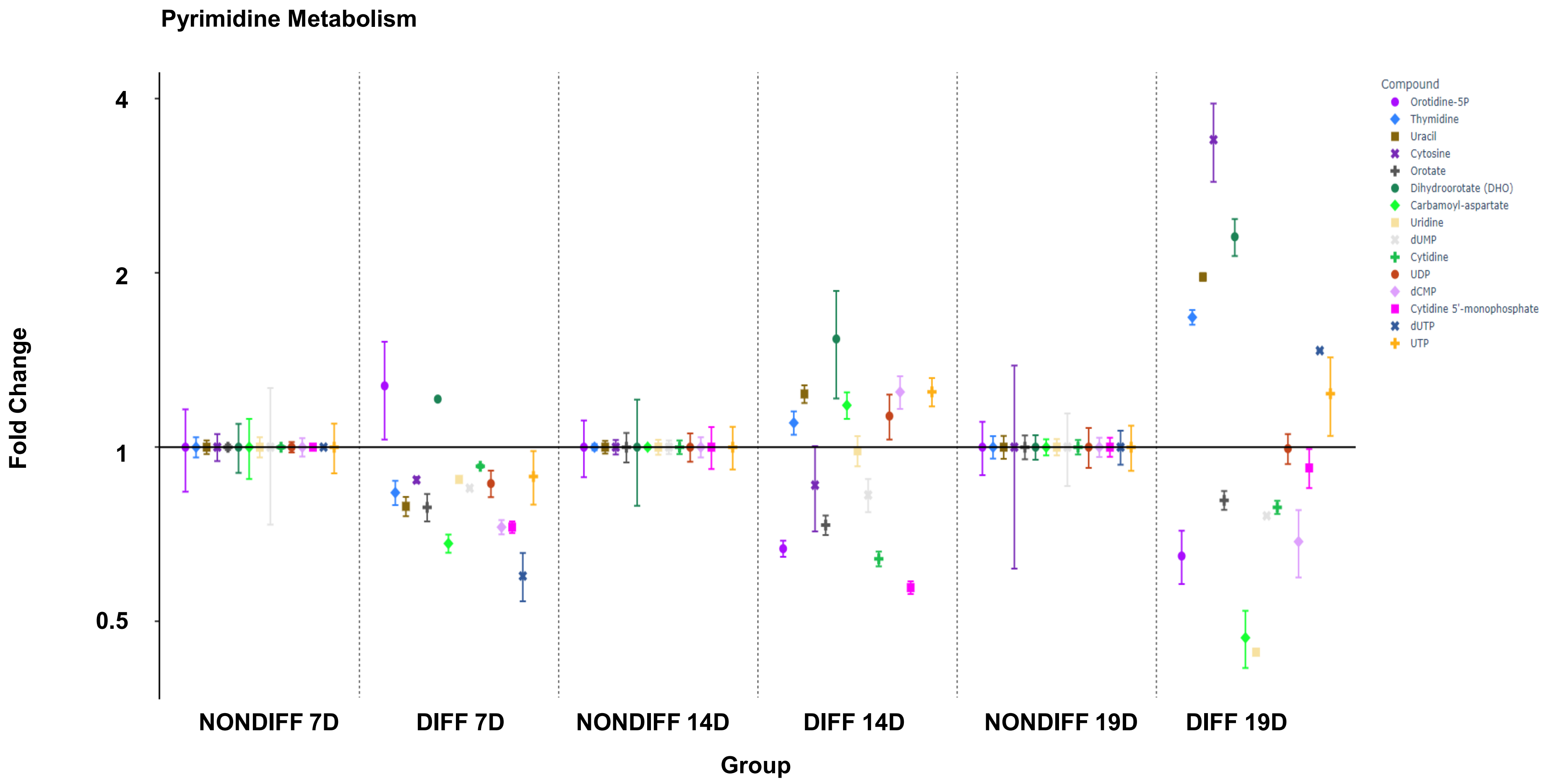


SFig5:De novo pyrimidine synthesis is altered during osteogenic differentiation. The panel presents relative fold changes in pyrimidine metabolites comparing non-differentiated and differentiated cells at multiple time points (NONDIFF_7d, DIFF_7d, NONDIFF_14d, DIFF_14d, NONDIFF_19d, DIFF_19d). Metabolites profiled include orotidine-5′-phosphate, dihydroorotate (DHO), carbamoyl-aspartate, orotate, uridine, UMP, cytidine, UDP, UTP and CTP. Non-differentiated cells remain largely near baseline, whereas differentiated cells show time-dependent increases in several intermediates (notably orotate, uridine, UDP/UTP and CTP) and decreases in others, indicating a shift in nucleotide biosynthetic flux. These data suggest that osteogenic differentiation is accompanied by reprogramming of pyrimidine metabolism to meet changing biosynthetic and energetic demands.

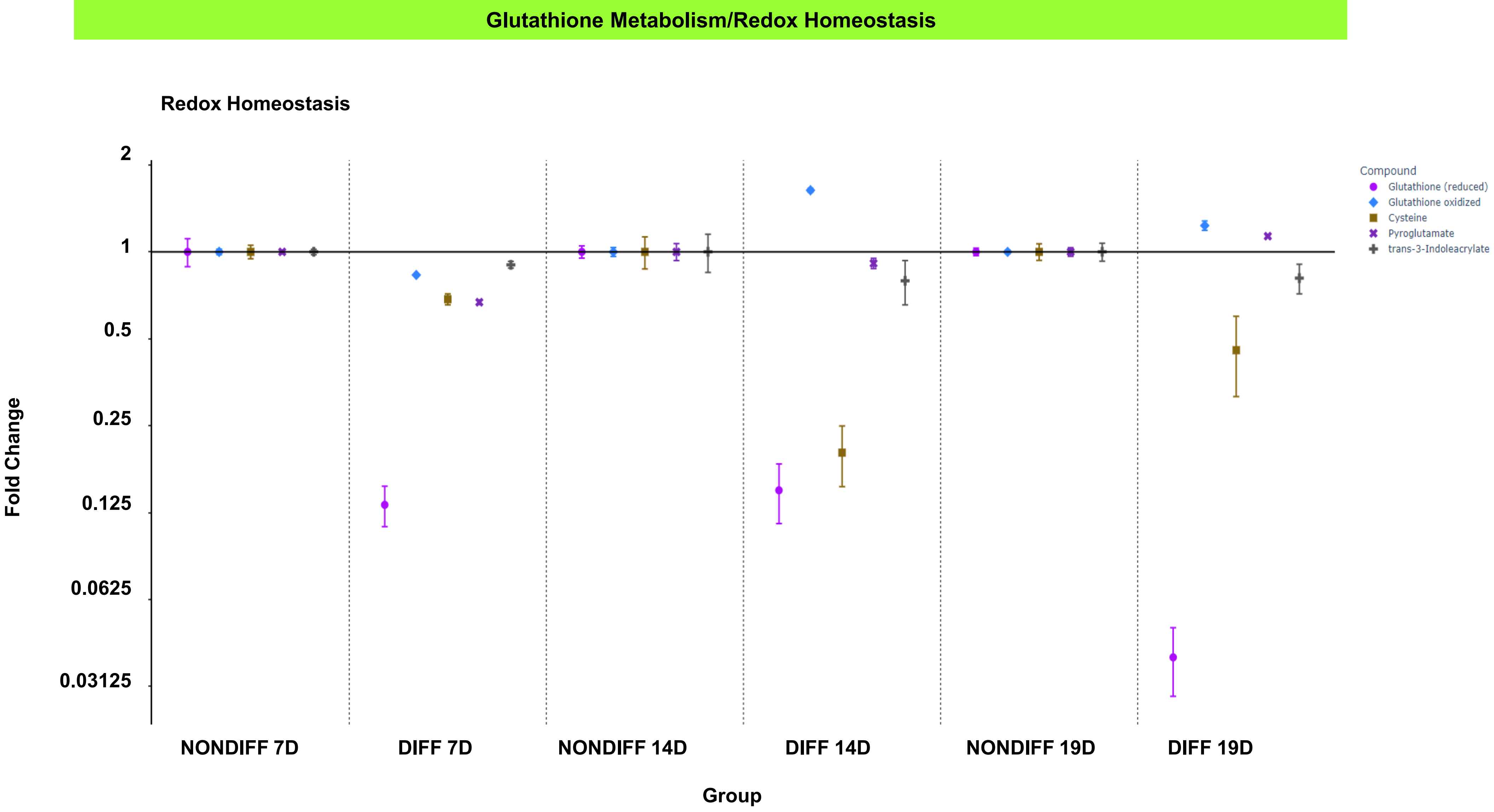


SFig6:Disruption of glutathione metabolism and redox homeostasis during osteogenic differentiation. The panel presents relative fold changes of redox-related metabolites across experimental groups (NONDIFF_7d, DIFF_7d, NONDIFF_14d, DIFF_14d, NONDIFF_19d, DIFF_19d). Metabolites include reduced and oxidized glutathione, cysteine, pyroglutamate, and trans-3-indoleacrylate. Non-differentiated samples remain largely near baseline, whereas differentiated cells display time-dependent alterations in glutathione intermediates, including a marked reduction in pyroglutamate and shifts in the balance of reduced versus oxidized glutathione at later time points. These data indicate impaired redox buffering capacity and increased oxidative stress during osteogenic differentiation.

| Gene | d7/d0 | ttest |
| --- | --- | --- |
| *Bnip3* | 7.6026719 | 0.011399 |
| *Bnip3l* | 1.4132369 | 0.071868 |
| *Bcl2l13* | 1.22 | NA |
| *Pink1* | 0.9 | NA |

SFig7: Data mined form a previously published data set showed Bnip3 expression is significantly upregulated during osteoblast differentiation of calvarial osteoblasts. Quantitative gene expression analysis comparing day 7 (d7) to day 0 (d0) of osteoblast differentiation shows a robust increase in Bnip3 expression (7.60-fold, p = 0.011), indicating activation of BNIP3 dependent pathways during osteoblast maturation. There was a slight trend towards an increase in Bnip3L and no significant differences in Bcl2l13 and Pink1 transcripts.

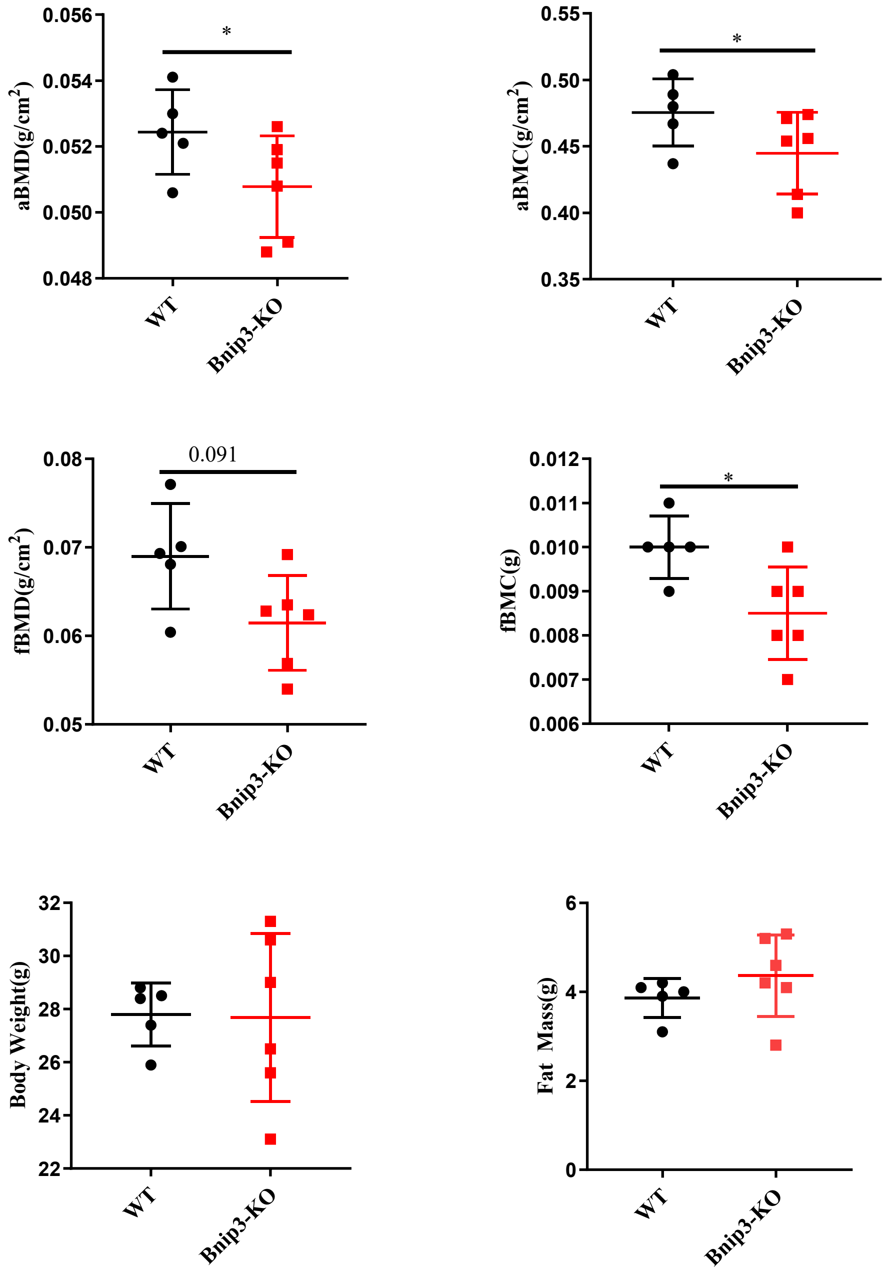


SFig8: Loss of BNIP3 decreases bone mineral density and content without significantly altering body composition. Dual-energy X-ray absorptiometry (DEXA) analysis demonstrates a significant reduction in areal bone mineral density (aBMD) and areal bone mineral content (aBMC) in Bnip3-KO mice compared with WT littermates ($p \leq 0.05$). Femoral bone mineral density (fBMD) shows a downward trend in Bnip3-KO mice ($p = 0.091$), while femoral bone mineral content (fBMC) is significantly reduced ($p \leq 0.05$). In contrast, total body weight and fat mass are not significantly different between genotypes, indicating that the skeletal deficits observed in Bnip3-KO mice are not secondary to changes in overall body composition. Each point represents an individual animal; bars indicate mean ± SEM.

### Supplemental Table 1

| Gene | Forward primer (5' to 3') | Reverse Primer (3' to 5') |
|---|---|---|
| *Alpl* | *CAT GTT CCT GGG AGA TGG TAT G* | *CTT GGA GAG GGC CAC AAA G* |
| *Atf4* | *TGC CGG TTT AAG TTG TGT GC* | *GGA TTT CGT GAA GAG CGC CAT* |
| *Bglap* | *CTG ACA AAG CCT TCA TGT CCA C* | *GCG CCG GAG TCT GTT CAC TA* |
| *Col1a1* | *CGT CTG GTT TGG AGA GAG CAT* | *GGT CAG CTG GAT AGC GAC ATC* |
| *Mfn2* | *CCT ACT GCT CCT TCT AAC CCA* | *CTG CCT CTC GAA TTC TGA AAC T* |
| *Hprt1* | *GCC TAA GAT GAG CGC AAG TTG* | *TAC TAG GCA GAT GGC CAC AGG* |
| *Bnip3* | *TCC TGG GTA GAA CTG CAC TTC* | *GCT GGG CAT CCA ACA GTA TTT* |
| *Bnip3l* | *ATG TCT CAC TTA GTC GAG CCG* | *CTC ATG CTG TGC ATC CAG GA* |
| *Bcl2l13* | *AGT GGA GAC TGC AGT CCA TG* | *CGT GCT CCT CCA GGT ACA TC* |
| *Runx2* | *CGA CAG TCC CAA CTT CCT GT* | *CGG TAA CCA CAG TCC CAT CT* |
| *Drp1* | *TGT TAC GGT TCC CTA AAC TTC AC* | *ATG CAC CAT TTC ATT TGT CAC G* |
| *Sp7* | *GAA GTT CAC CTG CCT GCT CTG T* | *CGT GGG TGC GCT GAT GT* |
| *Pink1* | *TTC TTC CGC CAG TCG GTA G* | *CTG CTT CTC CTC GAT CAG CC* |
